# Nanoparticles loaded with a CSF1R antagonist selectively depletes microglial cells and modulates inflammation in spinal cord injury

**DOI:** 10.1101/2024.11.21.624647

**Authors:** Jingjing Yang, Bernard Ucakar, Kevin Vanvarenberg, Alessio Malfanti, Anne des Rieux

## Abstract

Neuroinflammation is a principal event occurring after spinal cord injury (SCI). M1-like microglia are key players in the inflammatory response after injury. We hypothesize that the depletion of this microglia subtype would result in a more pro-resolutive environment, favorable to SCI repair. The colony-stimulating factor 1 receptor (CSF1R) antagonist PLX5622 has been used to deplete microglia in the central nervous system. Although PLX5622 can freely cross the blood-brain barrier after systemic administration, the low drug concentration within the SCI site hampers its effectiveness. Additionally, systemic administration of PLX5622 in the scope of SCI treatment can induce side effects due to off-target accumulation. In this study, we specifically depleted M1- like microglia by designing polymeric nanoparticles loaded with PLX5622 (PLX NPs) to locally treat spinal cord contusion. PLX NP was prepared using a microfluidic-assisted approach showing high encapsulation efficiency (approx. 84%), nanosized dimensions (100 nm), and batch-to-batch reproducibility. PLX NP displayed selective activity in depleting M1-like microglial cells in both resting and lipopolysaccharide (LPS)-activated mixed microglial cell models compared with the free drug counterparts while preserving non-targeted glial cells. Furthermore, locally administered PLX NP downregulated proinflammatory cytokines (e.g., TNF-α, IL-6, and IL-1β), increasing the M2/M1-like microglia ratio, thus reducing inflammation in a SCI contusion model. Our data support the hypothesis that local treatment with PLX NPs, a formulation with a high translational value, reduces neuroinflammation, with potential applications in SCI and central nervous system inflammatory diseases.

**Graphical abstract.:** The colony-stimulating factor 1 receptor (CSF1R) antagonist PLX5622 loaded in polymeric nanoparticles (PLX NP) is delivered for the treatment of neuroinflammation in spinal cord injury.

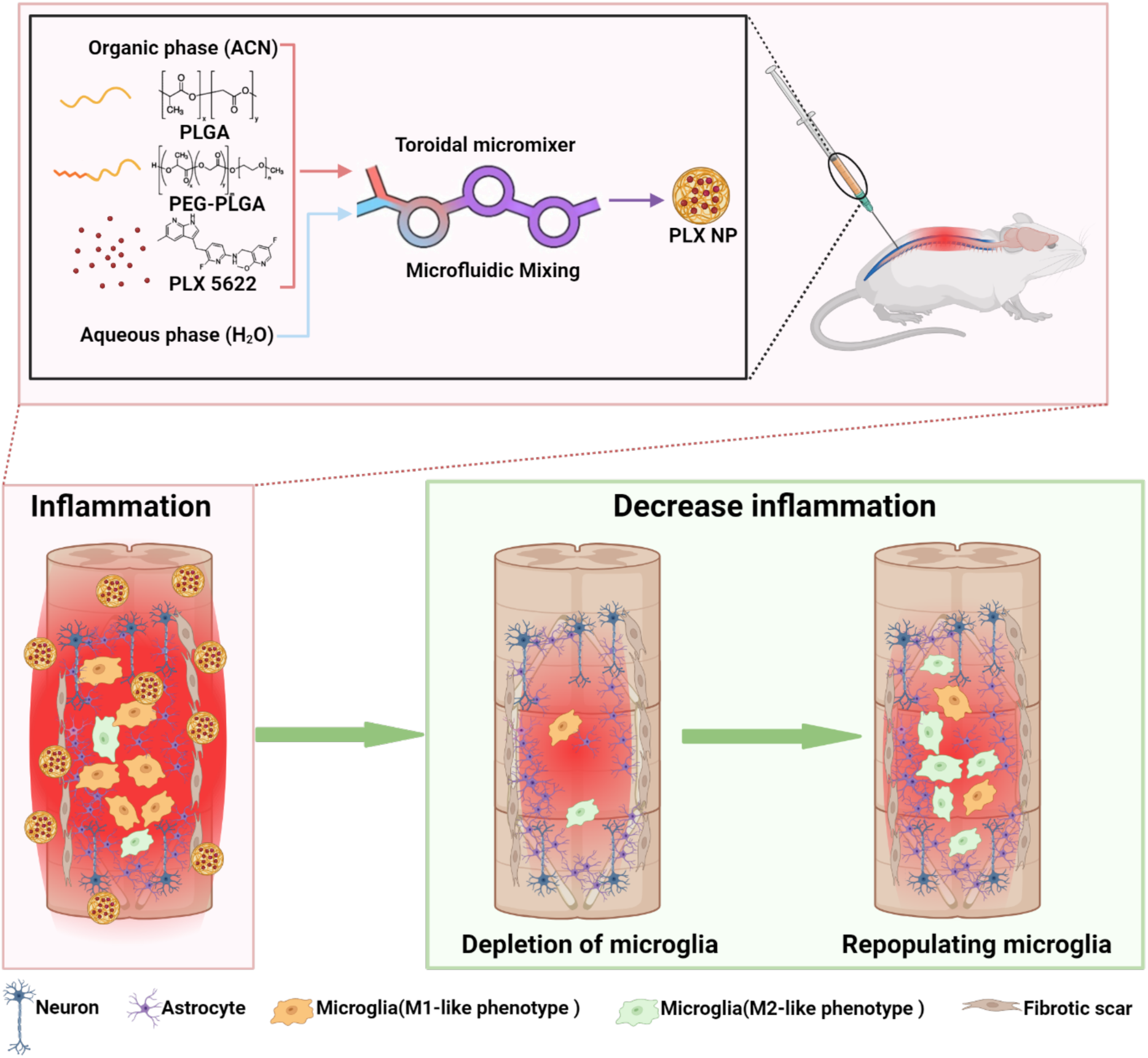

## 1 Introduction

Spinal cord injury (SCI) is one of the most critical public health concerns that results in permanent impairment of sensory, motor, and autonomic functions [1]. Every year, >500,000 new patients suffer from SCI mostly because of motor vehicle collisions and falls, leading to disability and high hospitalization costs [2]. Despite a greater understanding of the pathophysiology of SCI, neuroprotective and regenerative approaches allowing repair of the damaged spinal cord have had limited clinical efficiency to date [3].

Stem cell-based therapy is recognized as a promising strategy to treat SCI; however, its efficacy is greatly limited by low cell survival, notably impaired by local hyper-inflammation occurring after the lesion [4]. This phenomenon, also referred to as neuroinflammation, is one of the principal consequences of SCI and significantly contributes to the pathological progression from acute to chronic SCI [5]. Chronic inflammation after SCI leads to an unfavorable microenvironment for tissue repair and the viability of cells transplanted in the scope of cell therapy [6,7]. Thus, strategies aimed at resolving neuroinflammation would provide a better environment for spinal cord tissue recovery and for the success of other therapies [8]. Microglial cells are the main driver of this pathological condition and exhibit a gradient oscillating between two opposite phenotypes: a proinflammatory classically activated (M1-like) phenotype that supports the inhibition of cellular proliferation and a pro-resolutive alternatively activated (M2-like) phenotype that promotes inflammation resolution and tissue repair and homeostasis [9,10]. In the injured spinal cord, M1-like proinflammatory microglia are one of the main actors of chronic inflammation, whereas their shift toward a pro-resolutive phenotype (defined as M2-like) is impaired [5]. In this context, M1-like microglia could be a therapeutic target for SCI. Llyod *et al.* [11] showed that M1-like microglia death resulted in a resurgence of new pro-resolutive M2-like microglial cells and tissue repair [12]. Another study demonstrated that normalizing the SCI microenvironment led to the recovery of cognition and motor functions [13]. Among the recent strategies used to deplete microglial cells, the administration of colony-stimulating factor 1 receptor (CSF1R) antagonists has emerged as a promising approach for reducing chronic inflammation in several neurodisorders, including SCI, while improving neurological recovery [14,15]. Indeed, CSF1R antagonists can be used to modulate microglial populations in the central nervous system (CNS) when administrated orally [16]. However, the phagocytosis of macrophages and prolonged systemic administration of CSF1R antagonists have been associated with undesired off-target toxicity on peripheral immune cells, particularly those of lymphoid origin [17]. PLX5622 is a widely used CSF1R antagonist that has the lowest toxicity toward other glial cells like oligodendrocyte progenitor cells (OPCs) compared with other CSF1R antagonists, such as PLX3397 [18]. However, PLX5622 is poorly soluble in water and needs to be solubilized in DMSO before injection. This poor solubility, coupled with the risk of off-target side effects, significantly hinders its development for clinical use, particularly for local delivery to the injured spinal cord, where injection of DMSO is not permitted.

In this study, we hypothesized that the local delivery of PLX5622 will deplete M1-like microglial cells and restore a pro-resolutive microenvironment in SCI. To overcome the limitations of CSF1R antagonist therapy, we combined the encapsulation of PLX5622 in nanomedicines with local delivery by intrathecal administration. We selected poly(lactic-co-glycolic) (PLGA)-based nanoparticles (NPs) to encapsulate PLX5622 for its high translational value, as PLGAs are Food and Drug Administration (FDA)-approved, their formulation is easy and fast when produced by microfluidics, and does not require additional excipients such as surfactants. Microfluidics enables high encapsulation efficiency, high reproducibility, and easier scale-up [19]. Altogether, this ensures an easier translation into clinical settings. Here, we demonstrated for the first time that intrathecal injection of PLX5622 encapsulated in PLGA NPs induced a local depletion of M1-like microglia associated with the modulation of the SCI lesion toward a more pro-resolutive environment.

## 2 Experimental section

### 2.1 Materials

Poly(D, L-lactide-co-glycolide) (MW = 7,000–17,000) and poly(ethylene glycol) methyl ether-block-poly(lactide-co-glycolide) (PEG-PLGA) (PEG average Mn 5,000, PLGA Mn 7,000) were obtained from Sigma-Aldrich (Merck, Darmstadt, Germany). PLX5622 (PLX) (MW = 395.41) was obtained from Medkoo Biosciences, Inc. (North Carolina, USA). Dulbecco’s Modified Eagle’s Medium (DMEM), minimum essential medium (MEM), fetal bovine serum (FBS), N-2- hydroxyethylpiperazine-N-2-ethane sulfonic acid, penicillin–streptomycin (P/S), Hanks’ Balanced Salt Solution, 1,1′-dioctadecyl-3,3,3′,3′-tetramethylindodicarbocyanine 4- chlorobenzenesulfonate (DiD), PrestoBlue™ HS Cell Viability Reagent, and Trizol were obtained from Thermo-Fisher Scientific® (Bleiswijk, The Netherlands). LPS, bovine serum albumin (BSA), and Triton TM X-100 were purchased from Sigma-Aldrich (Merck, Darmstadt, Germany). O.C.T. compound, chloroform, isopropanol, and ethanol were purchased from VWR (Rosny-sous-Bois- cedex, France). The Go ScriptTM Reverse Transcription System and Go Taq qPCR master mix were purchased from Promega® Benelux BV (Leiden, The Netherlands). Tris-ethylenediaminetetraacetic acid buffer was obtained from Qiagen® (Hilden, Germany). Normal goat serum was obtained from Vector Laboratories (California, USA). Rat TNF-alpha ELISA Kit and Rat IL-10 ELISA Kit were purchased from Proteintech (Planegg-Martinsried, Germany).

### 2.2 Preparation of PLX-loaded polymeric NPs

PLGA (13.5 mg) and PEG-PLGA (1.5 mg) were dissolved in acetonitrile (ACN) (720 µL), and 0.75 mg of PLX5622 dissolved in DMSO was added to the polymer solution (30 µL). PLX-loaded NPs (PLX NPs) were produced by microfluidics using a NanoAssemblr Ignite^®^ Precision NanoSystems™ (Vancouver, BC, Canada) using a flow rate ratio of Milli-Q water (2.25 mL): acetonitrile of 3:1, a total volume of 3 mL, a total flow rate of 8 mL/min, a start waste of 0.45 mL, and an end waste of 0.15 mL. PLX NPs were then dialyzed against 1 L of Milli-Q water supplemented with 0.5% Pluronic® F-127 for 4 h at room temperature (RT) (Spectra/Por, MWCO: 6-8 kDa) and for 12 h in Milli-Q water to remove organic solvents and nonencapsulated PLX. Blank NPs were produced following the same protocol but with 30 µL of DMSO instead of PLX, whereas fluorescent PLX NPs were obtained by adding 7.5 µL of DiD (1mg/mL) to the organic phase. All formulations were sterile filtered on 0.22 μm and stored at 4°C until further use.

### 2.3 Characterization of PLX NPs

#### 2.3.1 PLX quantification in PLGA nanoparticles

Drug loading (DL) and encapsulation efficiency (EE) were evaluated using high-performance liquid chromatography (HPLC). PLX NPs (100 µL) were dissolved in 900 µL of ACN, and PLX was quantified by HPLC using a Shimadzu Prominence system (Shimadzu, Japan) equipped with a 150 × 4.6-mm column packed with a C18 column with a particle size of 5 µm (Macherey–Nagel, Germany). Water supplemented with ammonium acetate (0.02% m/v) (eluent A) and acetonitrile (eluent B) was used as the mobile phase in a gradient mode from 5% to 95% of eluent B at a flow rate of 1 mL/min. The detection wavelength was 280 nm, and the retention time was 11 min. DL and EE were calculated using the following formulas:

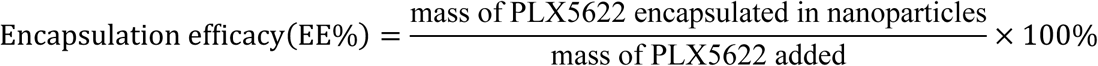

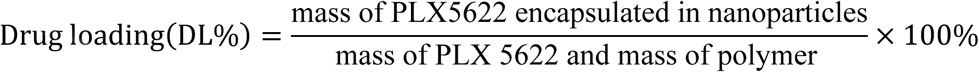

#### 2.3.2 Dynamic Light Scattering

The size, polydispersity index (PDI), and zeta-potential (ZP) of PLX NPs were measured in Milli-Q water using Malvern Zetasizer Nano ZS (Malvern Instruments) equipped with a red laser (λ = 633 nm) at a fixed angle of 173° at 25 °C. The PLX NP and Blank NP size and distribution were determined by using 50 µL of colloidal suspensions. The size values were reported as intensity.

#### 2.3.3 Transmission Electron Microscopy (TEM)

Ten μL of stock solution of PLX NP and Blank-NP in Milli-Q water were placed on a homemade carbon-coated copper grid, and the solvent was allowed to dry at room temperature. Following the time, samples were stained with 1% uranyl acetate in distilled water for 2 min at room temperature to provide for negative staining. TEM analyses were performed using a FEI Tecnai G2 microscope (Hillsboro, OR, USA). NP size analysis was performed with ImageJ Software version 1.52a (National Institutes of Health, USA). The average size of the PLX NP and Blank NP was calculated by analyzing 5-20 individual particles in 10 different measurements with ImageJ software.

#### 2.3.4 PLX in vitro release

PLX release was studied in complete cell culture medium (including 10% FBS) and in cerebrospinal fluid (CSF) obtained from Cliniques Universitaire Saint-Luc (ethics committee approval 2007/10SEP/233, the patients included signed an internal regulatory document, stating that residual samples used for diagnostic procedures can be used for retrospective academic studies, without any additional informed consent.) between January 2018 and January2022, and the samples comes from classified as symptomatic controls which were defined as patients in whom a thorough neurological examination, combined with biological and MRI imaging data, did not reveal any underlying neurological disease. PLX NP were suspended in medium or CSF (10% (v/v) of PLX NP) and incubated at 37°C under agitation (120 rpm). At different time points, samples were centrifuged (n=3), the supernatant removed and the amount of PLX quantified by HPLC, as described above, after the pellet dissolution in 10% water, 89% ACN and 1% DMSO.

### 2.4 Depletion of microglial cells in vitro

Primary mixed glial cells (MGCs) were obtained from neonatal Wistar rats (1–2 days old, De Duve Institute Animal Facility, UCLouvain), as previously described [20]. Briefly, the brains were removed aseptically, and the cerebellum was separated. After the asportation of the meninges, brains were disrupted mechanically and digested for 45 min (papain in MEM (40 µg/mL)). Next, digested tissues were centrifuged at 160 g 4℃ for 5 min and resuspended, triturated slowly with 18- and 23-gauge needles, and filtered on a 40-µm cell strainer (Greiner Bio-One, Frickenhausen, Germany). The cells were then resuspended in growth medium (DMEM containing glucose, L-glutamine, pyruvate, 10% FBS, 1% penicillin, and 2.5% HEPE) and seeded in 24-well plates (10^6^ cells/well) and 96-well plates (1.68×10^5^ cells/well). The cells were further incubated for 8 days at 37°C in a 5% CO_2_ incubator. After 8 days, MGCs were treated with different concentrations of PLX (in 0.1% DMSO), Blank NP, and PLX NP (0.01–40 µM of PLX). The same experiment was performed on LPS-activated MGCs. MGCs were treated with LPS (100 ng/mL) for 2 h and incubated for 72 h with PLX, Blank NP, and PLX NP (4 µM and 20 µM of PLX) diluted in growth medium supplemented with fresh LPS (100 ng/mL). The effect of treatment on MGCs was analyzed using a cell viability assay, immunofluorescence, and for the LPS experiment by RT-qPCR and ELISA (as per supplier instructions).

### 2.5 Cell viability

The impact of PLX NP on MGC viability was examined using a PrestoBlue™ HS Cell Viability Reagent (Thermo-Fisher Scientific^®^, Bleiswijk, The Netherlands) following supplier instructions. After incubation with the different treatments, the PrestoBlue™ solution was added to each well and incubated for 10 min. Cell metabolic activity was measured using a fluorimeter (Molecular Devices Spectramax^®^ iD5, California, USA) at λ_ex_= 560 nm and λ_em_= 600 nm. Cell culture media (background), untreated MGCs (100% viability), and MGCs treated with 0.1% Triton X-100 (0% viability) were used as controls.

### 2.6 Immunofluorescence

Microglial cell depletion was analyzed by immunofluorescence against Iba-1. MGCs were washed once with PBS before fixation (15 min, paraformaldehyde (PFA) 4%) and then washed three times with PBS. The cells were incubated with Iba-1 primary antibodies (Table S1) diluted in 10% (v/v) normal goat serum in PBS for 16 h at 4 ℃. The cells were washed three times with 0.01% PBS-Tween 20 and then incubated for 3 h with secondary antibodies **(Table S1)** and 4′,6-diamidino-2- phenylindole (DAPI) (1:1,000) diluted in PBS containing 5% normal goat serum. The cells were washed three times with 0.01% PBS-Tween 20 and stored in PBS. Images of whole wells were acquired and analyzed using an ImageXpress Pico Automated Cell Imaging System (IX-PICO, California, USA) (4 ×, 10 ×, and 20 × objective). The cells were detected by DAPI staining, positive cells were detected using a fluorescence intensity classification method and summed. Results were expressed as the number of total cells and positive cells.

### 2.7 RT-qPCR

Total RNA from cells was extracted using the Trizol reagent, as previously described [21]. One μg of total RNA was used to synthesize cDNA using the GoScript Reverse Transcription System. qPCR was performed using a StepOnePlus instrument and software (Applied Biosystems), as previously described [21]. Primer sequences for qPCR are listed in **Table S2**.

### 2.8 Impact of PLX local administration on spinal cord LPS-induced inflammation in a mouse model

The experimental protocol was approved by the UCLouvain local ethics committee (2022/UCL/MD/056). LPS (300 μg/kg, i.p.) was administered to induce CNS inflammation [21]. Four hours after LPS injection, PLX NPs (10 μL of 10 μg PLX) were injected (20 µL/min) using a 25-μL syringe and Remote Infuse/Withdraw Pump 11 Elite Nanomite Programmable Syringe Pump (Massachusetts, USA) in the intrathecal space between the groove of L5 and L6 vertebrae (tail flick) (n = 6) [22]. Mice were treated once a day for 3 days, and spinal cords were retrieved on day 4, 24 h after the last injection. Mice injected with only PBS intrathecal injection (i.t.) (vehicle; no LPS) and mice injected with LPS (intraperitoneal (i.p.) injection) followed by i.t. injection of PLX solubilized in 5% DMSO were used as negative controls. Mice injected with LPS followed by i.t. injection of PBS were used as positive controls. Spinal cords were then collected and stored at -80°C for RT-qPCR analysis or fixed in 4% PFA by transcardiac perfusion for immunofluorescence staining.

### 2.9 Mouse spinal cord immunofluorescence staining

Spinal cords were incubated in 20% sucrose for 24 h followed by incubation in 30% sucrose for 24 h and embedded in Tissue-Tek^®^ O.C.T.™ (Sakura, AJ Alphen aan den Rijn, The Netherlands). Cryosections of 10 μm were mounted on SuperFrost® slides (Thermo-Fisher Scientific, Brussels Belgium) and stored at −20°C until staining. Slides were allowed to dry at RT overnight before staining. Sections were blocked with PBS containing 10% normal goat serum, 2% BSA, and 0.2% Triton X-100 for 1 h and incubated with primary antibodies overnight at 4°C (Table S1). The sections were washed with PBS-Tween 80 (0.1%) four times before incubation with the appropriate secondary antibodies (Table S2) and DAPI for 2 h at RT. Immunofluorescent staining was visualized and digitalized using a Pannoramic 250 FlashIII scanner (3DHistech) at x20 magnification. Scanned slides were then analyzed using the image analysis tool Qupath 0.4.3. On each slide, tissue sections were automatically delimited at low magnification. Tissue areas were detected using a thresholding classification method with moderate (1.38µm/px) and max channels, and stained pixels in the lesion were detected (high resolution [x20])) using a thresholding classification method and summed. Results were expressed as a percentage of the stained area within the entire tissue.

### 2.10 Whole blood flow cytometry

To assess the potential systemic toxicity of PLX NPs, the impact of the treatment on circulating macrophages was analyzed using FACS in mouse whole blood. At the end of the experiment, 0.5 mL of blood was sampled from each mouse in the retrobulbar venous plexus and collected in heparinized tubes. For each antibody and for each single-color control (used to set compensation), aliquots of 100 μL of whole blood were transferred into 12 × 75-mm tubes. Whole blood samples were used as unstained controls for background setup. αF4/80, αCD86, and αCD206 primary antibodies (Table S1) (1:100 in Hanks’ Balanced Salt buffer) were added to the samples and incubated for 1 h at 4°C under agitation protected from light. The samples were washed with 2 mL of staining buffer, and the cells were resuspended in 500 μL of staining buffer. Analysis was performed using the Spectral Flow Cytometry (Cytek Aurora) **(Figure S10)**.

### 2.11 Impact of local PLX administration on spinal cord inflammation in a mouse spinal cord contusion model

Animal experiments were approved by the UCLouvain local ethics committee (2023/UCL/MD/46). Swiss female mice (5–6 weeks old) were treated with Temgesic (0.05 μg/g) and Carprofen (5 μg/g) by subcutaneous injection 30 min before surgery. The mice were anesthetized using a rodent anesthesia system with vaporized isoflurane (2%) before laminectomy at T8–T10 and spinal cord exposure. SCI was performed using a contusion of 50 kDynes (dwell time of 1 s) applied at the T9 level using a Horizon Spinal cord Impactor (WPI, Friedberg, Germany). The mice were then administered with Baytril (enrofloxacin, 2.5 mg/kg s.c.) and Temgesic (0.05 μg/g s.c.) once/day, and the bladder was emptied twice per day after surgery [23]. The following day, the mice were randomly divided into five groups: healthy mice treated with vehicle (PBS) by intrathecal injection (Control); SCI treated with vehicle (PBS) by i.t. injection (Veh); SCI treated with PLX solution in 5 % DMSO by i.p. injection (PLX i.p.); SCI treated with PLX solution in 5 % DMSO by i.t. injection (PLX i.t.); and SCI treated with PLX NP by i.t. injection (PLX NP i.t.) (n=10). Mouse body weight was recorded every day. Treatments were administered once a day for 3 days following SCI [24]. Spinal cords were collected 24 h after the last injection and stored at −80°C for RT-qPCR analysis (n=5) or fixed in 4% PFA by transcardiac perfusion for immunofluorescence analysis (n=5) [23]. Immunofluorescent staining was visualized and digitalized using a Pannoramic 250 FlashIII scanner (3DHistech) at x20 magnification. Scanned slides were then analyzed using the image analysis tool Author version 2017.2 (Visiopharm, Hørsholm, DNK). On each slide, tissue sections were automatically delimited at low magnification. Lesions were then manually delineated and stained pixels in the lesion and concentric circles were detected (high resolution [x20])) using a thresholding classification method and summed. Results were expressed as a percentage of stained area within the analyzed region. *Statistical analysis*: Statistical analyses were performed using one-way analysis of variance (ANOVA), two-way ANOVA, or nested one-way ANOVA using Prism 9 (GraphPad Software, San Diego, CA). Results are expressed as means ± standard deviations or means ± standard errors of the mean (as mentioned in figure legends) and were considered significant if *p* < 0.05. The number of replicates is indicated in each figure legend.

## 3 Results

### 3.1 Characterization of polymeric NPs loaded with PLX (PLX NPs)

We developed a highly efficient, reproducible, and scalable microfluidic-assisted approach (Ignite®, Precision NanoSystems™) to produce PLX NPs **(Figure 1a**) with high encapsulation efficiency (EE= 84%) and drug loading (DL=4.35%). PLX NP and Blank NP showed a round/ovoidal shape and presented a similar average hydrodynamic diameter (98 ± 2 nm and 100 ± 3, respectively) confirmed by dynamic light scattering (DLS) and TEM, a low polydispersity index (0.16 ± 0.01 and 0.17 ± 0.02, respectively) and a negative surface charge (−34 ± 2 mV and -33 ± 2 mV, respectively) (**Figure 1b-e**). Importantly, our methodology ensures strong reproducibility with negligible differences among the three batches (**Figure S1a-b**). PLX release studies were performed in media mimicking in vitro and in vivo conditions using DMEM medium supplemented with 10% FBS and in human-derived Cerebral Spinal Fluid (**Figure 1f**). Results show an initial burst drug release after 10 h of incubation of PLX NP in the relevant media followed by a sustained release of PLX along the time.

**Figure 1.**
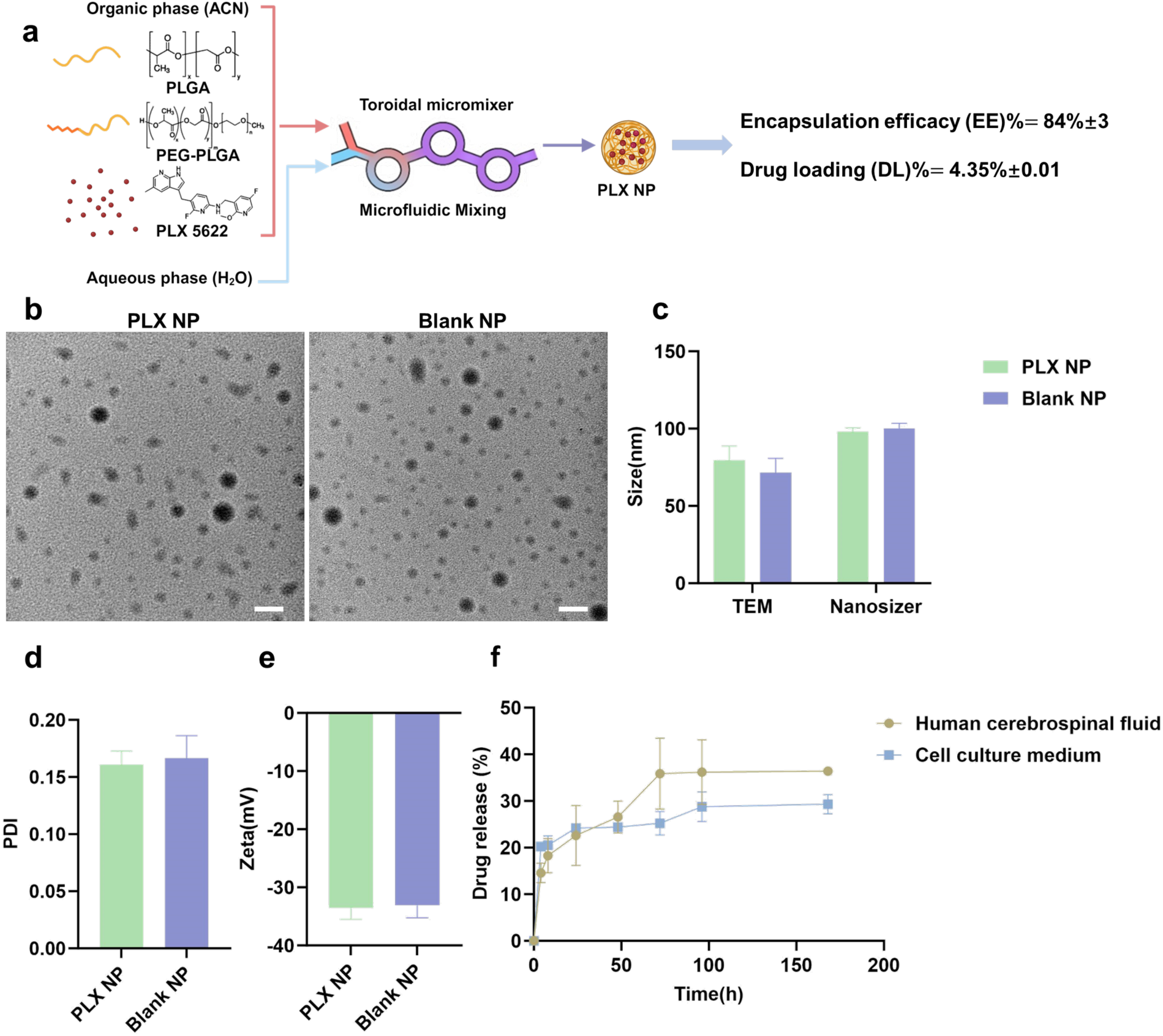
PLX NP characterization. (a) Formulation of PLX NP by Microfluidics using the Ignite® instrument and relative EE% and DL% of PLX NP. Data: mean ± SD (N=3, n=3/group). PLX NP and Blank NP size by TEM (b) and DLS (c), PDI (d), and Zeta potential (e). Scale bar: 100 nm. (f) PLX release profile from NP in cell culture media and human-derived Cerebral Spinal Fluid.

### 3.2 PLX NP selective depletion of microglial cells in vitro

We hypothesized that PLX NPs may specifically deplete microglial cells in a Mix Glial Cells (MGCs) model. The effects of PLX NPs on microglial cells were evaluated by immunofluorescence against Iba-1, a microglia marker, after incubation with MGC for different times and at different concentrations.

When MGCs were incubated for 72 h with PLX NPs, a significant decrease in the Iba-1-positive population was observed, starting at 0.5 µM of PLX **(Figure 2a)**. From 2 µM, >90% of the microglial population was depleted. The other cell types were not affected by the treatment up to 8 µM where a slight decrease in the cell number was observed; however, it was never >20%. Treatment with free PLX depleted 90% of the Iba-1-positive population at 0.5 µM; however, it was slightly more toxic to other cell types than PLX NPs because 25% of their population was depleted **(Figure 2b)**. Blank NPs had no impact on MGCs **(Figure S2)**. When comparing the different conditions, MGCs needed to be treated with at least 4 µM of PLX NPs to obtain the same level of depletion observed with PLX **(Figure 2c and Figure S3)**. The different treatments had no impact on the general MGC viability **(Figure S4)**. Only the highest PLX concentrations (20 and 40 µM) induced a slight decrease in the global MGC viability, likely because of microglia and OPC depletion. MGCs needed to be treated for at least 24 h to see a significant effect of PLX, whether as a solution or encapsulated in NPs **(Figure 2d and Figure S5)**.

**Figure 2.**
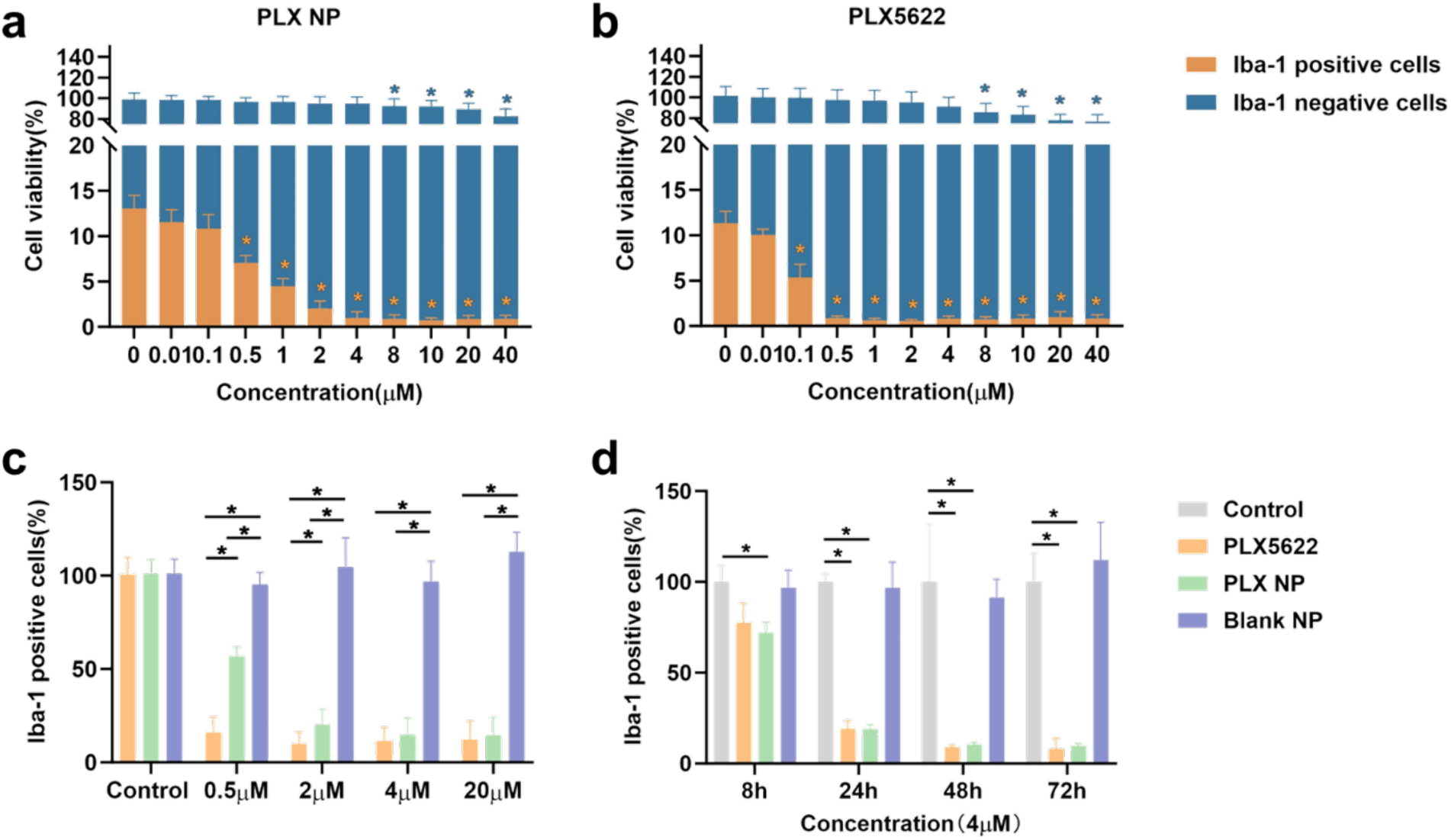
Impact of PLX NP on selective depletion of microglial cells in vitro. MGC cells were incubated with different concentrations of (a) PLX, (b) PLX NP and Blank NP for 72 h (c) (N=3, n=8). MGC cells were incubated with 4 μM of PLX, PLX NP and Blank NP for 8 h, 24 h 48 h and 72 h (d) (N=3, n=6). Then cells were stained for Iba- 1 (microglial cells) and DAPI (nuclei) and images were acquired with an IX-PICO. Analysis was done on whole wells using IX-PICO software. Data are expressed as the mean ± SD. Statistical analysis was done using a Nested one-way ANOVA or two-way ANOVA (\**p* < 0.05).

### 3.3 Off-target toxicity of PLX NPs on astrocytes and oligodendrocytes

MGC cultures contain microglia, astrocytes, OPCs, and oligodendrocytes [25]. Because our objective was to specifically deplete microglial cells, the off-target toxicity of PLX NPs on other cells was examined. Treatment with PLX or PLX NPs had no impact on astrocytes (GFAP+) **(Figure 3a)**. In contrast, a decrease in oligodendrocytes (Olig2+) was observed following treatment with both PLX and PLX NPs. However, PLX encapsulation limited the depletion of cells from the oligodendrocyte lineage, particularly at higher PLX concentrations **(Figure 3b)**.

**Figure 3.**
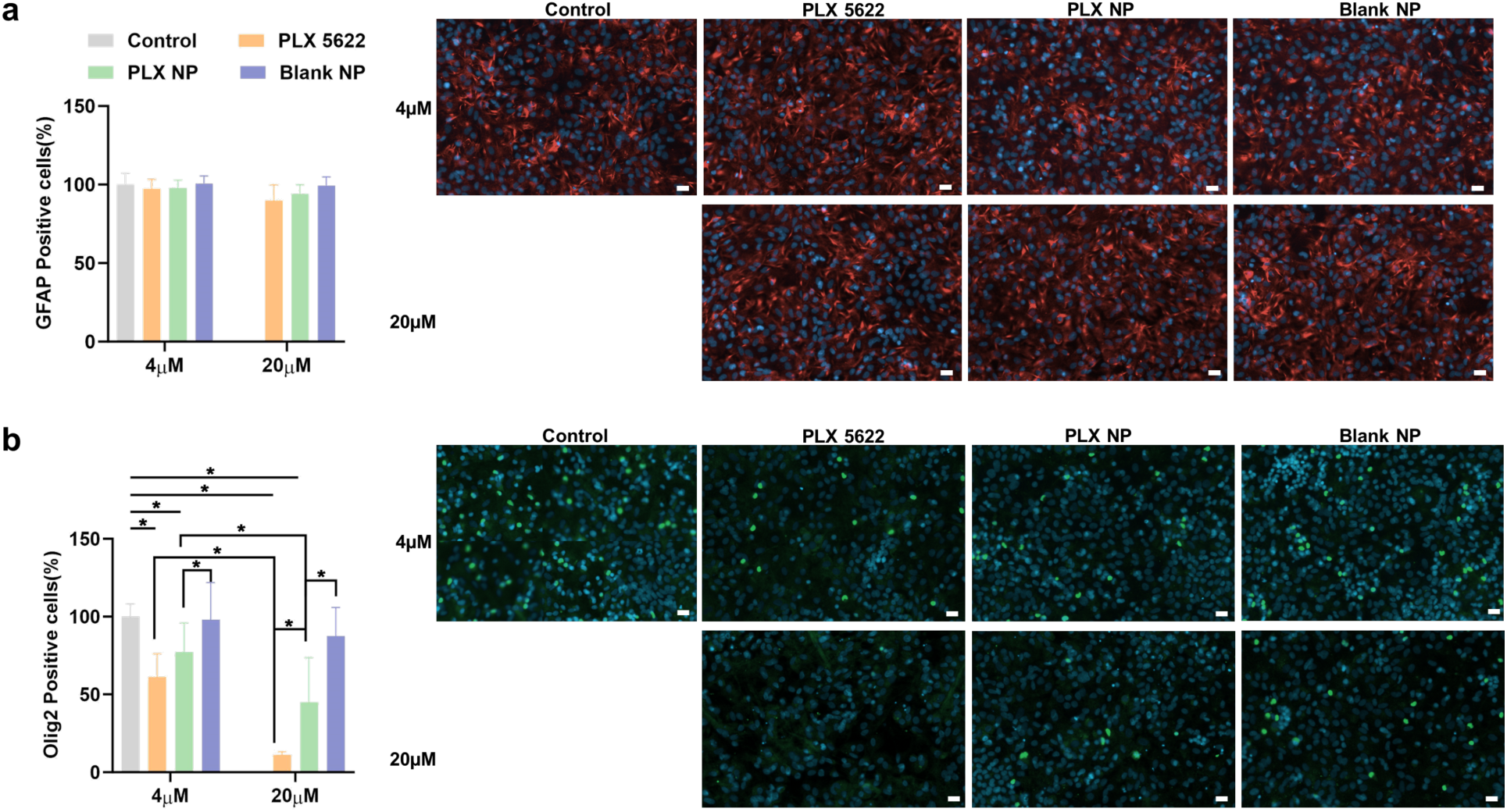
Impact of PLX NP on astrocytes and cells from the oligodendrocyte lineage. MGC cells were incubated with 4μM or 20 μM of PLX, PLX NP and Blank NP for 72h (N=3, n=7). Then off target toxicity was studied on astrocytes (a, stained with GFAP, green) and on oligodendrocytes (b, Olig-2, green) cells. Images were acquired with an IX-PICO instrument. Analysis was performed on whole wells using IX-PICO software. Data are expressed as the mean ± SD. Statistical analysis was done using two-way ANOVA (*p < 0.05). Scale bar: 20 μm, DAPI-blue, GFAP- red, Olig2-green.

### 3.4 Influence of MGC activation on PLX NP efficiency

Because PLX NPs were designed to act in a neuroinflammatory environment, the next step was to assess how MGC activation would impact PLX NP efficiency. MGCs were activated with LPS and treated with PLX NPs for different incubation times. Microglial cells were quantified by immunofluorescence against Iba-1. As observed for the previous experiment, at least 48 h was required to see significant microglial depletion. Activated microglial cells seemed less affected by PLX and PLX NPs than resting cells because only 50% of the population was depleted by treatment with 4 µM of PLX or PLX NPs compared with 90% in resting cells **(Figure 4a, b)**.

**Figure 4.**
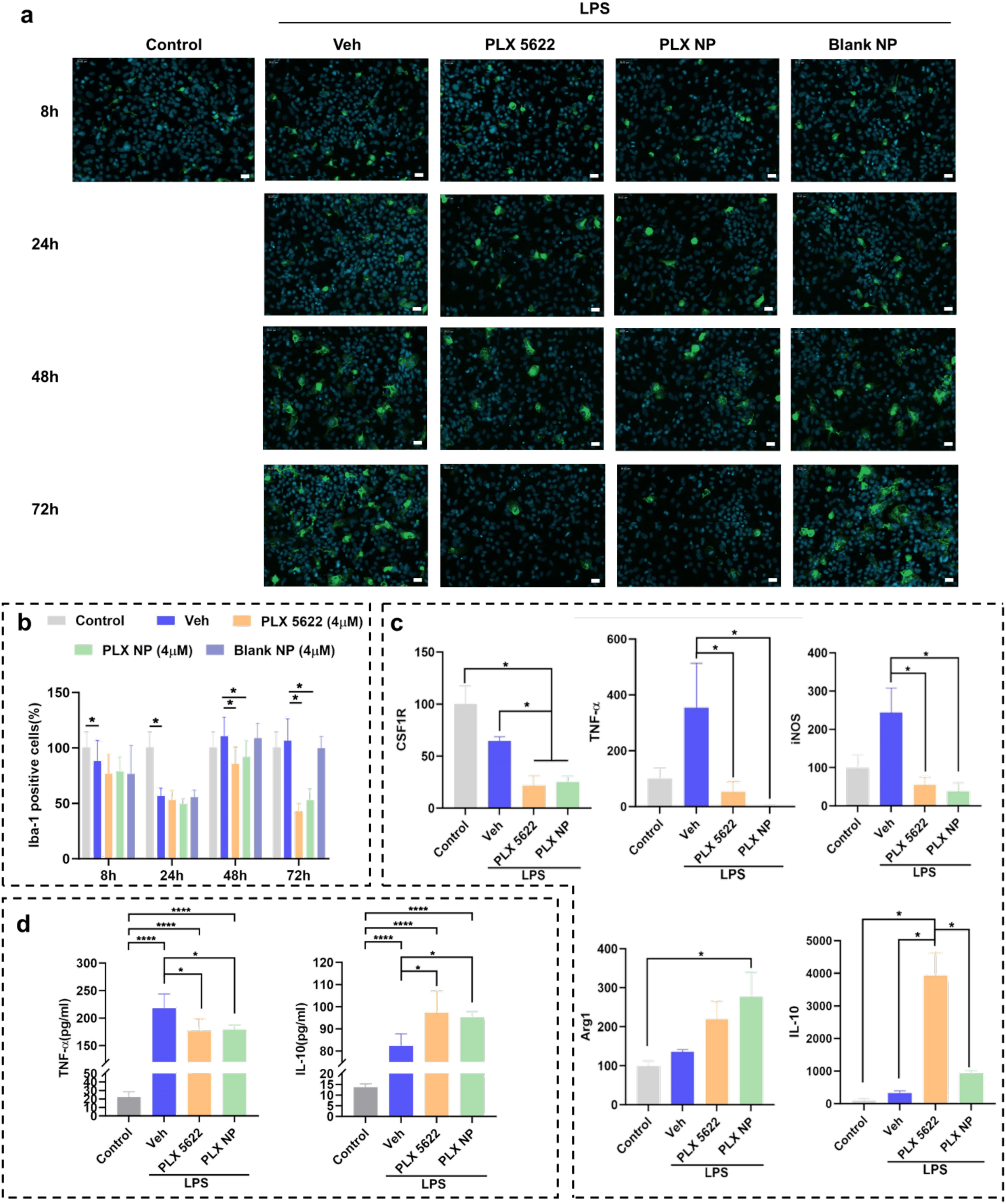
PLX NP efficiency on LPS-activated MGC and inflammatory cytokine gene expression. LPS-activated MGC (Veh) were incubated with 4μM of PLX, PLX NP and Blank NP for 72h (N=3, n=5-6). (a) Cells were stained for Iba-1 (microglia cells) and DAPI (nuclei) and images were acquired with an IX-PICO. (b) Analysis was done on whole wells using IX-PICO software. Data are expressed as the mean ± SD. Statistical analysis was done using two-way ANOVA. Scale bar: 20 μm, DAPI-blue, Iba-1-green; (c) CSF1R, iNOS, TNF-α, Arg1 and IL-10 gene expression was analyzed by RT-qPCR. (d) ELISA analyzed TNF-α and IL-10 expression in cell supernatant. Data are expressed as the mean ± SD and are normalized versus non-activated non-treated cells (Control). Statistical analysis was done using one-way ANOVA (\**p* < 0.05).

Next, we examined how the gene expression of inflammatory cytokines (both pro- and anti- inflammatory) was affected by MGC activation by LPS after PLX NP treatment. Treatment of LPS-activated MGCs with PLX NP significantly reduced CSF1R, iNOS, and TNF-α gene expression (proinflammatory markers) but increased Arg1 expression (pro-resolutive marker) compared with those in cells treated with the vehicle **(Figure 4c)**. Only PLX induced a significant increase in IL-10 gene expression (pro-resolutive marker). The decrease of TNF-α and the increase of IL-10 in the supernatants following cell treatment with PLX were confirmed by ELISA **(Figure 4d)**.

### 3.5 PLX NP uptake by resting and activated MGCs

Next, we evaluated whether PLX NPs could be endocytosed by resting and activated MGCs and whether a preferential cell-specific uptake occurred. DiD-labeled PLX NPs (**Figure S1c**) were endocytosed by both activated and resting MGCs in a time-dependent manner **(Figure 5a-b and Figure S6)**. PLX NPs were preferentially taken up by microglial cells (Iba-1+) compared with astrocytes (GFAP+) and oligodendrocytes (Olig2+) without significant differences between the resting and activated states. Furthermore, we observed a higher fluorescent signal associated with NPs in astrocytes than in oligodendrocytes.

**Figure 5.**
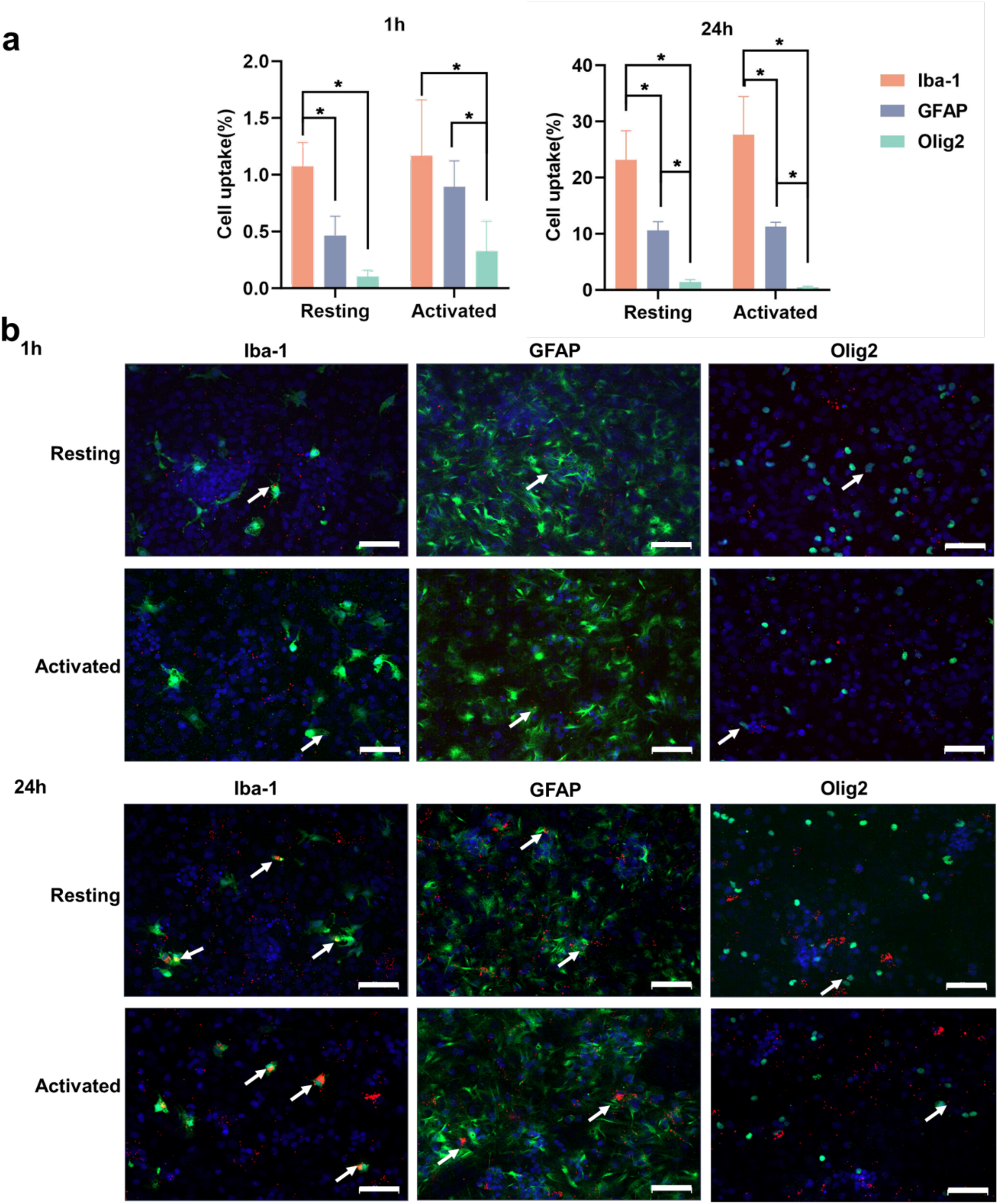
PLX NP uptake in resting and activated MGC. MGC activated by LPS (Activated) or not (Resting) were incubated with 4 μM of PLX NP- DiD for 1 h and 24 h (N=3, n=4). (a) Cell uptake was quantified using IX-PICO software. (b) Microglial cells were stained for Iba-1, astrocytes for GFAP, oligodendrocyte lineage cells for Olig2 and nuclei with DAPI. Data are expressed as the mean ± SD. Statistical analysis was done using two-way ANOVA (\**p* < 0.05). Scale bar: 50 μm, DAPI- blue, Iba-1-green, GFAP- green, Olig2- green, PLX NP- DiD- red.

### 3.6 Influence of PLX NPs on spinal cord microglial cells in an inflamed mouse model

We then explored the impact of PLX NPs on microglia and neuroinflammation in an LPS-induced inflammation model. Mice were treated with 300 µg/kg LPS by i.p. injection and then with an intrathecal injection of PLX NPs every day for 3 days **(Figure 6a, Figure S7a)**. The impact of PLX NPs on the spinal cord was evaluated using RT-qPCR and immunofluorescence. Furthermore, the CD206/CD86 (pro-resolutive/proinflammatory macrophages) ratio was analyzed in mouse blood using FACS. The treatments had no impact on mouse weight **(Figure S7b)**.

**Figure 6.**
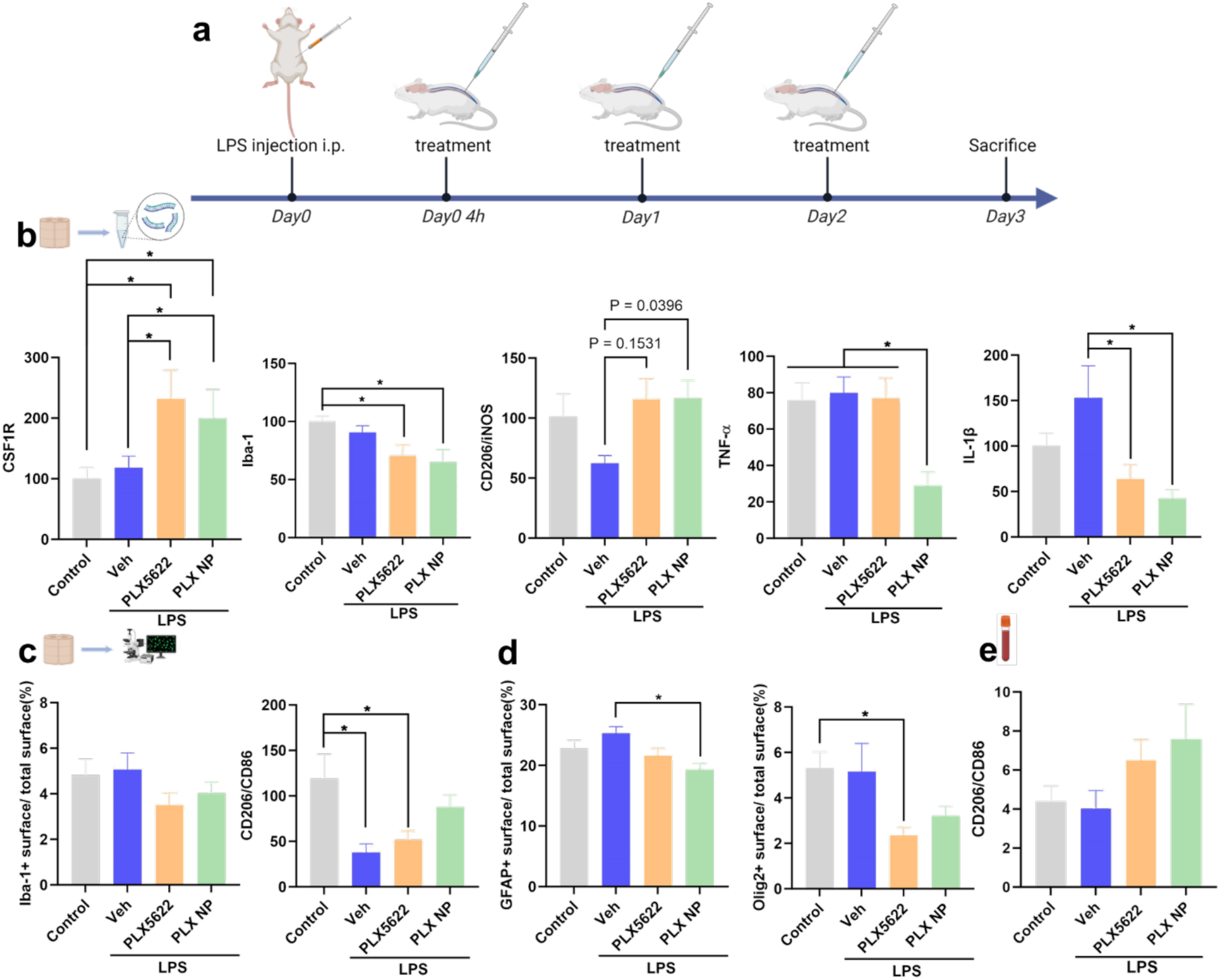
Impact of PLX NP on spinal cord gene expression in a LPS activated model. (a) Illustration of the experimental setting. (b) At day 3, mRNA was extracted from spinal cord and gene expression was analyzed by RT- qPCR. (c) Spinal cords were immunostained for Iba-1 (microglia cells), CD86 (M1-like phenotype microglia), CD206 (M2-like phenotype microglia), (d) GFAP (astrocytes), olig2 (oligodendrocyte lineage) and DAPI (nuclei) and images were acquired with a slide scanner. Analysis of the whole sections was done using Qu-Path. (e) Whole blood CD206/CD86 ratio was measured by FACS. Data are expressed as the mean ± SEM. Statistical analysis was done using One-way ANOVA (\**p* < 0.05) (n=6).

LPS injection did not affect Iba-1 and TNF-α gene expression, whereas it significantly decreased the CD206/iNOS ratio (pro-resolutive/proinflammatory macrophages) and increased IL-1β gene expression **(Figure 6b)**. The injection of PLX NPs significantly decreased Iba-1, TNF-α, and IL- 1β gene expression, whereas PLX decreased Iba-1 and IL-1β expression. Only PLX NPs significantly decreased TNF-α expression and significantly increased the CD206/iNOS ratio.

LPS injection had no effects on the percentage of Iba-1-positive staining in the spinal cord of LPS-treated mice, whereas it significantly decreased the CD206/CD86 ratio **(Figure 6c and Figure S7c)**. However, only the injection of PLX NPs allowed the CD206/CD86 ratio to increase to a level similar to that in the positive control (non-LPS-activated, PBS i.t. injection).

Finally, when analyzing whole blood for the ratio between CD206+ and CD86+ macrophages, no significant difference was observed; however, the ratio tended to increase for PLX- and PLX NP-treated animals **(Figure 6e)**.

The toxicity of PLX NPs on the astrocyte and oligodendrocyte populations was also evaluated. Injection of LPS did not increase GFAP- or olig2-positive staining; however, administration of PLX NPs significantly decreased GFAP-positive staining, whereas only the injection of PLX significantly depleted cells of the oligodendrocyte lineage **(Figure 6d and Figure S7d)**.

### 3.7 Impact of local PLX NP administration on a contusion model of SCI

Finally, we assessed our hypothesis that PLX NPs modulate the local microenvironment in a spinal cord contusion model. PLX NPs were administered intrathecally 1 day after contusion, and their effect on the spinal cord was evaluated using RT-qPCR and immunofluorescence **(Figure 7a)**. PLX and vehicle (PBS) were used as negative controls, whereas intraperitoneal injection of PLX (PLX i.p.) was used as a comparison with what is usually performed in the literature [24]. The treatments had no impact on mouse weight **(Figure S8a)**. One mouse died during surgery. and the PLX i.p. group lost 20% of its mice after the first day of treatment, suggesting that PLX i.p. injection was toxic.

**Figure 7.**
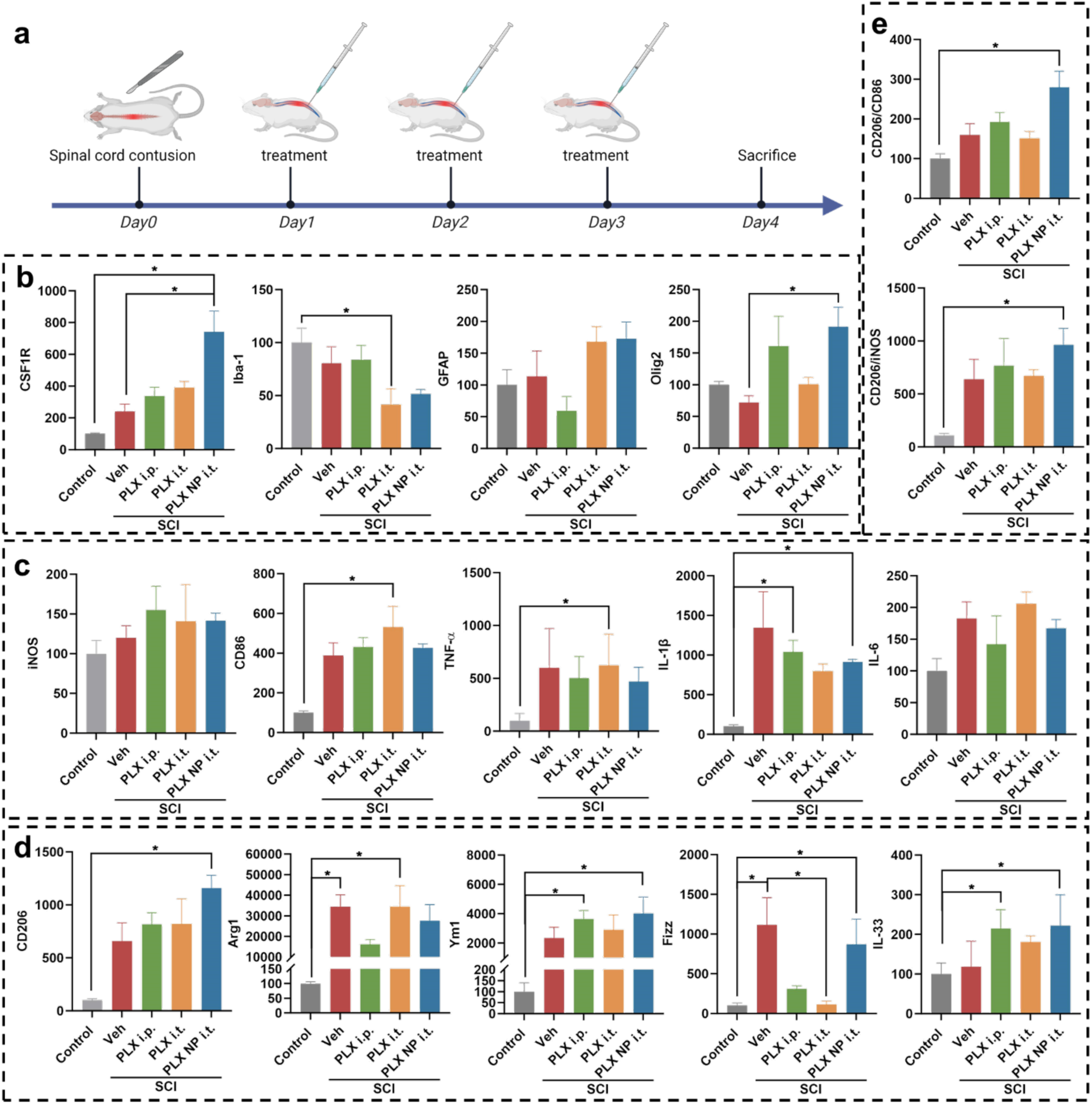
PLX NP influence on spinal cord gene expression after SCI. (a) Illustration of the experimental setting used for this experiment. Gene expression of (b) CSF1R, Iba-1, GFAP and Olig2 was analyzed by RT-qPCR. Similarly, gene expression of M1-like phenotype and pro-inflammatory markers (c) iNOS, CD86, TNF-α, IL-1β and IL-6 and M2-like phenotype and anti-inflammatory markers (d) CD206, Arg1, Ym1, Fizz, and IL-33 were measured by RT- qPCR. M2/M1 ratios (e) CD206/CD86 and CD206/iNOS ratio were calculated. Data are expressed as the mean ± SEM. Statistical analysis was done using One-way ANOVA (\**p* < 0.05) (n=5).

Intrathecal injection of PLX NPs after contusion induced a significant increase in CSF1R gene expression compared with every other condition **(Figure 7b)**. Iba-1 expression decreased when the mice were treated with PLX and PLX NPs. The expression of the olig2 gene increased upon PLX NP injection; however, GFAP expression was unaffected. I.p. injection of PLX tended to decrease GFAP and increase olig2 although this decrease was insignificant.

PLX NPs had no statistically significant effect on the gene expression of proinflammatory markers, such as iNOS, CD86, TNF-α, IL-6, and IL-1β (slight decrease but insignificant) **(Figure 7c)**. Treatment with PLX NPs increased the expression of pro-resolutive markers, such as CD206, Ym1, Fizz, and IL-33, and increased the CD206/iNOS and CD206/CD86 ratios **(Figure 7d and 7e, respectively)**. Intraperitoneal injection of PLX also increased Ym1 and IL-33 but decreased Fizz gene expression. When analyzing macrophages in the whole blood and in the spleen, none of the treatments displayed systemic toxicity **(Figure S8b and Figure S8c)**.

These results were confirmed by immunofluorescence at the lesion site. PLX NPs increased the ratio of CD206/CD86-positive staining but had no significant effect on Iba-1-, CD86-, and CD206- positive staining or GFAP, NeuN, and olig2 **(Figure 8 and Figure S9)**.

**Figure 8.**
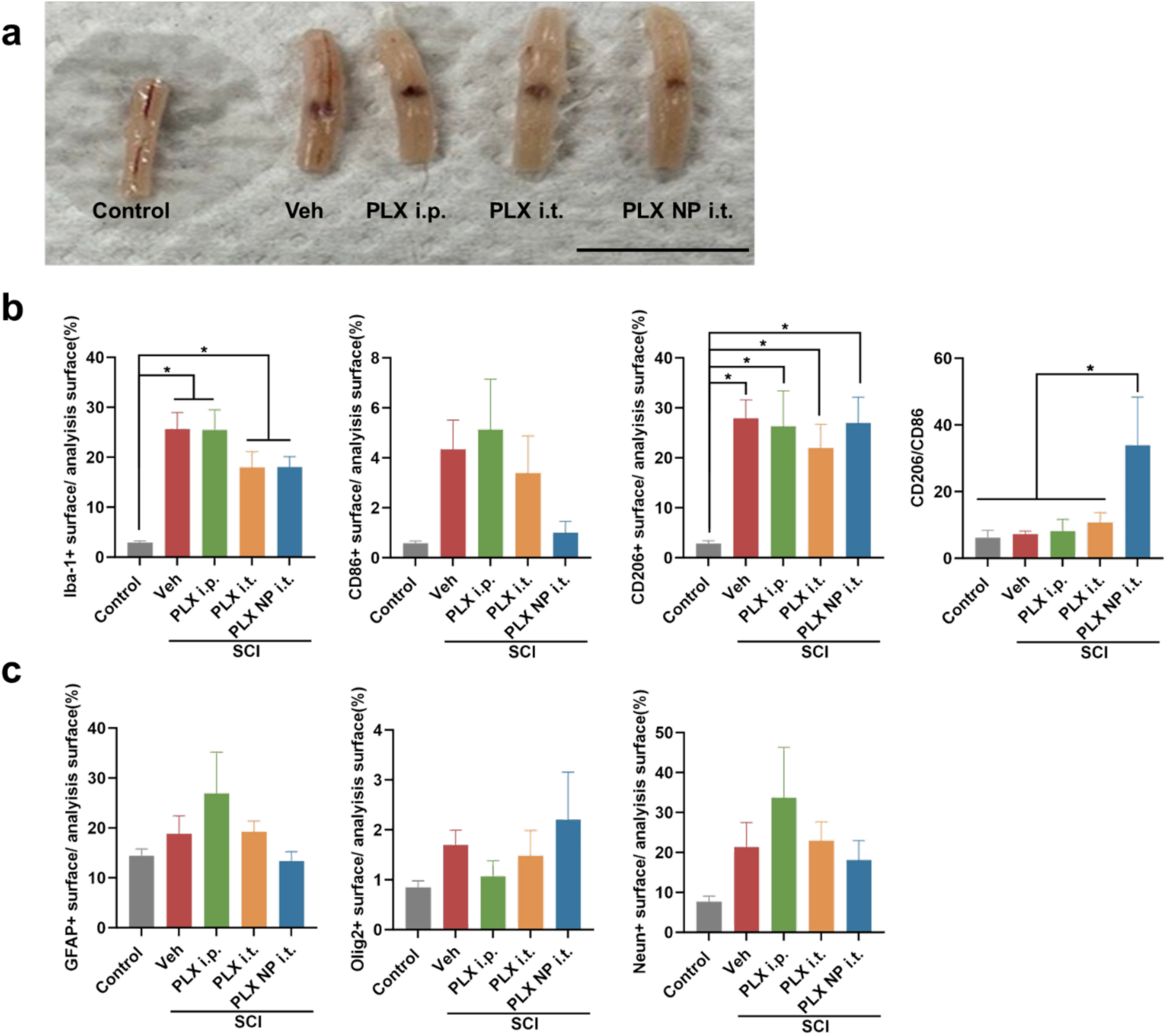
Evaluation of PLX NP effect on spinal cord tissue after SCI. Spinal cords (a) were immunostained at the lesion site (Scale bar: 1 cm) for (b) Iba-1 (microglia cells), CD86 (M1-like phenotype microglia), CD206 (M2-like phenotype microglia), and (c) GFAP (astrocytes), Olig2 (oligodendrocyte lineage cells), NeuN (neuronal cells) and DAPI and Images were acquired with a slide scanner. Analysis of the lesion site was done using VisioPharm. Data are expressed as the mean ± SEM. Statistical analysis was done using One-way ANOVA (\**p* < 0.05) (n=5).

## 4 Discussion

This study established a novel approach for the local treatment of SCI using a nanomedicine that selectively depletes microglial cells and modulates inflammation. M1-like microglia emerged as a key player in generating a proinflammatory microenvironment, hampering the healing of the spinal cord injured area. Therefore, the local depletion of this cell subtype represents an interesting strategy for treating SCI while avoiding systemic side effects. The design of the PLX NPs enhances the effect of the CSF1R antagonist PLX5622 preferentially in the microglial cells because of their high phagocytosis rate while avoiding the toxicity in other cells, such as astrocytes, oligodendrocytes, and OPCs. Moreover, the depletion of M1-like microglia has been reported to contribute to restore homeostasis and thus to resolve chronic inflammation and support neurological recovery [12,26]. Previously, PLX5622 has been administered systemically with undesired off-target effects on peripheral immune cells [27,28]. Therefore, we developed a new efficient system for controlled and sustained delivery of CSF1R antagonists to create a favorable microenvironment for the recovery of SCI. We selected PLGA as a matrix for the PLX NPs and microfluidics as a production method to provide a high translation value to our nanomedicines. To the best of our knowledge, this is the first report of a PLX-type CSF1R encapsulation in a fit-for-purpose nanomedicine locally administered for SCI repair.

As far as we know, we are the first to produce nanomedicines loaded with PLX5622. Encapsulation in PLGA NP provides the well-known advantages of solubilization (no need to co-inject DMSO, toxic for neuronal cells and not allowed in patients receiving local administration in the spinal cord) [29–31], protection from degradation, and controlled drug release. Fostering translation to the clinics, we produced PLX NPs using a microfluidic-assisted approach ensuring high batch-to-batch reproducibility and scalability. Indeed, bigger instruments can be used with minimal adjustments to produce clinical batches (up to 17 L/h), as is the case for mRNA-loaded LNP [32]. Microfluidics allowed the production of NP with a narrower size distribution than those obtained with traditional mechanical methods. It is also faster and requires fewer steps and manipulation. We demonstrated that our delivery system allows a biphasic release of PLX in relevant media such as cell culture and human-derived cerebral spinal fluid. We hypothesize that the initial fast release might be related to the PLX that is absorbed on the outer shell of the PLGA nanocarrier and therefore is released faster. Then, the release slowed, which can be classically observed, especially for lipophilic drugs, as their release is often influenced by their ability to bind to proteins in the medium and transition from the PLGA core to the solution.

We showed that PLX NPs can efficiently deplete microglia in a MGC *in vitro* model while limiting off-target effects on other cells, such as astrocytes and oligodendrocytes. The PLX concentration required to obtain the same effect compared with free PLX was higher. It is to be expected since an encapsulated drug is always less accessible than a free molecule *in vitro* and can enter cells via different mechanisms. PLX NPs efficiently and selectively depleted microglia after only 24 h of incubation. Although the mechanism by which PLX NPs deplete microglial cells remains unknown, we speculate this to be more related to the amount of NPs taken up by the cells than PLX release because >20% of the microglial cells endocytosed NPs after 24 h while release was quite slow, at least in PBS. Studies have indicated that CSF1R is localized in the nucleus and that the full-length CSF1R migrates to the nucleus upon CSF1 stimulation [33], where it could be susceptible to PLX released from internalized NPs.

To further explore the therapeutic potential of our formulation, we examined the efficacy of PLX NPs in LPS-activated MGCs to better mimic the SCI-inflamed microenvironment. We observed a reduction of PLX efficiency regardless of its encapsulation. This has not been reported so far.

Considering that the percentage of microglia that endocytosed PLX NPs was the same regardless of whether the MGCs were activated or not, there is no clear explanation for this observation. We saw a 35% decrease in CSF1R expression in LPS-activated MGCs (not statistically significant however) and a reduction of approximately 75% in CSF1R when the cells were incubated with PLX or PLX NPs. A decrease in CSF1R expression in the presence of LPS has already been reported [34]. It was attributed to a crosstalk between Toll-like receptor and CSF-1 signaling pathways where LPS induces a pro-TNF-α-converting enzyme that cleaves CSF1R, resulting in ectodomain shedding [35]. CSF1R is then removed from the cell surface, and CSF1 signaling is blocked. This could explain why PLX seems less efficient in the presence of LPS. Treatment with PLX NPs significantly decreased the expression of the proinflammatory markers iNOS and TNF- α while increasing the expression of the anti-inflammatory makers Arg1 and IL-10. This is consistent with the results of an earlier study that observed the same effect when treating microglial cells with another CSF1R antagonist (Ki20227) [36]. The lower uptake of PLX NPs by other cells, such as astrocytes and most oligodendrocytes/OPCs, is advantageous to limit off-target effects and toxicity toward remyelinating cells. PLX5622 has been selected for its higher selectivity toward microglial cells and lower affinity for OPCs [18]; however, encapsulation in NPs further reinforces its targeted effect while reducing off-target effects.

Microglial depletion in the scope of brain diseases has been used to reset microglia to a homeostatic phenotype; however, to the best of our knowledge, the therapeutic potential of PLX NPs for the local treatment of SCI has not been explored yet. To evaluate the feasibility of our approach, we first investigated the impact of PLX NPs on a model of neuroinflammed spinal cord using LPS. The intrathecal administration of free PLX or PLX NPs led to comparable microglia depletion. However, only PLX NPs increased the M2/M1 ratio (both at the gene and protein expression levels) and decreased proinflammatory cytokine gene expression (TNF-α and IL-1β) compared with the LPS control group. This agrees with what has been reported in other models. For instance, PLX5622 more specifically depleted CD86 (M1-like) macrophages than CD206 (M2-like)- expressing ones and reduced the expression of proinflammatory cytokines (TNF-α and IL-1β) in a partial sciatic nerve ligation model [37]. In a mouse contusion traumatic brain injury model, microglial depletion by PLX5622 was followed by a subsequent repopulation of neuroprotective microglia [38]. However, treatment with PLX5622 is not always successful because Vichaya et al. [39] reported that it did not abrogate the inducible expression of proinflammatory cytokines in the brain in response to LPS and even exacerbated it for some cytokines.

Because we observed a slight decrease in GFAP+ cells (approximately 24%), it is not excluded that a slight depletion of astrocytes could participate in the reduction of proinflammatory markers observed when mice were treated with PLX NPs. Astrocytes are known to participate in SCI neuroinflammation and glial cell scarring [40,41]. Therefore, this can be considered an additional beneficial effect of PLX NPs. Witcher et al. showed that microglial depletion could attenuate astrocyte inflammatory and reactive signatures in a traumatic brain injury model [42]. PLX encapsulation in NPs also limits its toxicity on cells from the oligodendrocyte lineage. This, associated with an activated microglia decrease, would preserve remyelination potential [43].

Finally, the ability of PLX NPs to modulate the inflammatory response following SCI was evaluated in a mouse contusion model. Although PLX NP and PLX administration had no impact on blood and spleen macrophages, intraperitoneal administration of PLX resulted in the death of 20% of the mice after a single administration. This confirms the interest and safety of the local administration of PLX NPs for SCI. Interestingly, treatment by intrathecal administration of PLX NPs increased CSF1R gene expression in the LPS and contusion models. Even if the timing of our experiment was short, this could reflect the initiation of the repopulation occurring by the proliferation of the surviving microglia [44]. Here we opted to treat the mice 24h after the spinal cord contusion as we aimed to limit the intensity of the inflammation before its installation. However, it has been shown that inflammation could be beneficial to some extent in the acute phase of SCI due to debris clearing [45]. For further experiments where we will explore the impact of PLX NPs on spinal cord repair, starting PLX NP injections 3 days post-injury might be more appropriate.

As observed with the LPS model, treatment with PLX NPs depleted activated microglia and tended to decrease astrocyte activation while sparring cells from the oligodendrocyte lineage. SCI results in the loss of oligodendrocytes and thus demyelination, compromising efficient conduction and the long-term health of axons [46]. Protection of oligodendrocytes and remyelination are thus key factors for preventing axonal degeneration and restoring function following SCI. As observed in the LPS model, intrathecal injection of PLX NPs decreased the gene expression of some proinflammatory markers; however, in this study, its effect was more marked on pro-resolutive markers and on the M2/M1 ratio. In the scope of SCI, it is beneficial for the repair process to reduce inflammation and to induce the polarization of microglial cells toward a M2-like phenotype at the earlier stage [47] particularly because the secondary injury mechanisms initiate within 24 h after SCI, leading to exacerbation of tissue injury [48].

## 4. Conclusion

Progress in nanomedicine technologies and pharmacological target selection are crucial to ameliorate the outcomes of patients with SCI. In this study, we demonstrated that the local administration of PLX NPs in the subdural space efficiently modulated the injured microenvironment by decreasing M1-like microglia and simultaneously increasing M2-like microglia in a murine SCI model. Intrathecal injection of PLX NPs could deplete microglia and increase the M2/M1 ratio at the lesion site, while limiting off-target effects and systemic toxicity, providing a pro-resolutive microenvironment for spinal cord repair and potentially for cell therapy.

Our approach has the advantages (i) to use FDA-approved biomaterials that facilitate translation into clinical settings; (ii) to employ a production method for PLX NPs that is highly reproducible and with high scaling-up potential; (iii) to modulate the injured microenvironment, opening the possibility to exploit combinational approaches, such as cell therapy; and (iv) to be easily applied by surgeons in the injured area while reducing systemic side effects. Our work provides a strategy against neuroinflammation by resetting microglia to a homeostatic phenotype, with potential applications for SCI and other CNS inflammation diseases.

## CRediT authorship contribution statement

**Jingjing Yang:** Writing – original draft, Visualization, Validation, Methodology, Investigation, Formal analysis, Data curation, Conceptualization. ***Bernard Ucakar*:** Methodology, Formal analysis. ***Kevin Vanvarenberg*:** Methodology, Formal analysis, Data curation. ***Alessio Malfanti*:** Writing – review & editing, Resources, Investigation, Supervision, Conceptualization, Funding acquisition. ***Anne des Rieux*:** Writing – review & editing, Resources, Investigation, Supervision, Funding acquisition, Conceptualization.

## Data Availability

The data that support the findings of this study are available from the corresponding author upon reasonable request.

## Acknowledgments

J.Y. was supported by a China Scholarship Council (CSC) fellowship. A.M. was supported by the Marie Skłodowska-Curie Actions for an Individual European Fellowship under the European Union’s Horizon 2020 research and innovation program (grant agreement no. 887609). AdR is supported by Fonds de la Recherche Scientifique—Fonds National de la Recherche Scientifique (FRS-FNRS) (grant agreement no. 40000747) (Belgium). The authors would like to thank the Fondation Louvain for funding the acquisition of the Ignite instrument and Prof. Sophie Lucas for access to the ImageXpress Pico Automated Cell Imaging System. The authors would like to thank Dr. Bouzin (UCLouvain, IREC Imaging platform (2IP)) for her help with the image quantification and Prof. van Pesch (UCLouvain, IoNS) for the CSF. We thank Dr. Federico Caicci for the support with the TEM instrument and analysis settings.

## Supplementary data

Supplementary Information is available.

## References

[1] N.B. Jain, G.D. Ayers, E.N. Peterson, M.B. Harris, L. Morse, K.C. O’Connor, E. Garshick, Traumatic spinal cord injury in the United States, 1993-2012, JAMA. 313(22) (2015) 2236-43. doi: 10.1001/jama.2015.6250.

[2] M. Karsy, G. Hawryluk, Modern Medical Management of Spinal Cord Injury, Curr Neurol Neurosci Rep. 19(9) (2019) 65. doi: 10.1007/s11910-019-0984-1.

[3] T.H. Hutson, S. Di Giovanni, The translational landscape in spinal cord injury: focus on neuroplasticity and regeneration, Nat Rev Neurol. 15(12) (2019) 732–745. doi: 10.1038/s41582-019-0280-3.

[4] D. Liu, G. Lu, B. Shi, H. Ni, J. Wang, Y. Qiu, L. Yang, Z. Zhu, X. Yi, X. Du, B. Shi, ROS-Scavenging Hydrogels Synergize with Neural Stem Cells to Enhance Spinal Cord Injury Repair via Regulating Microenvironment and Facilitating Nerve Regeneration, Adv Healthc Mater. 12(18) (2023) e2300123. doi: 10.1002/adhm.202300123.

[5] J.C. Gensel, B. Zhang, Macrophage activation and its role in repair and pathology after spinal cord injury, Brain Res. 1619 (2015), 1619:1–11. doi: 10.1016/j.brainres.2014.12.045.

[6] H. Xue, B. Ran, J. Li, G. Wang, B. Chen, H. Mao, Bone marrow mesenchymal stem cell exosomes-derived microRNA-216a-5p on locomotor performance, neuronal injury, and microglia inflammation in spinal cord injury, Front Cell Dev Biol. 11 (2023) 1227440. doi: 10.3389/fcell.2023.1227440.

[7] V. Neirinckx, D. Cantinieaux, C. Coste, B. Rogister, R. Franzen, S, Concise review: Spinal cord injuries: how could adult mesenchymal and neural crest stem cells take up the challenge? Stem Cells. 32(4) (2014) 829–43. doi: 10.1002/stem.1579.

[8] X. Freyermuth-Trujillo, J.J. Segura-Uribe, H. Salgado-Ceballos, C.E. Orozco-Barrios, A. Coyoy Salgado, Inflammation: A Target for Treatment in Spinal Cord Injury, Cells. 11(17) (2022) 2692. doi: 10.3390/cells11172692.

[9] Z.S. Gao, C.J. Zhang, N. Xia, H. Tian, D.Y. Li, J.Q. Lin, X.F. Mei, C. Wu, Berberine-loaded M2 macrophage-derived exosomes for spinal cord injury therapy, Acta Biomater. 126 (2021) 211–223. doi: 10.1016/j.actbio.2021.03.018.

[10] Y. Tan, T. Lai, Y. Li, Q. Tang, W. Zhang, Q. Liu, S. Wu, X. Peng, X. Sui, F. Reggiori, X. Jiang, Q. Chen, C. Wang, An oil-in-gel type of organohydrogel loaded with methylprednisolone for the treatment of secondary injuries following spinal cord traumas, J Control Release. 374 (2024) 505–524. doi: 10.1016/j.jconrel.2024.08.033.

[11] A.F. Lloyd, C.L. Davies, R.K. Holloway, Y. Labrak, G. Ireland, D. Carradori, A. Dillenburg, E. Borger, D. Soong, J.C. Richardson, T. Kuhlmann, A. Williams, J.W. Pollard, A. des Rieux, J. Priller, V.E. Miron, Central nervous system regeneration is driven by microglia necroptosis and repopulation. Nat Neurosci. 22(7) (2019) 1046–1052. doi: 10.1038/s41593-019-0418-z.

[12] Y. Huang, Z. Xu, S. Xiong, F. Sun, G. Qin, G. Hu, J. Wang, L. Zhao, Y.X. Liang, T. Wu, Z. Lu, M.S. Humayun, K.F. So, Y. Pan, N. Li, T.F. Yuan, Y. Rao, B. Peng, Repopulated microglia are solely derived from the proliferation of residual microglia after acute depletion, Nat Neurosci. 21(4) (2018) 530–540. doi: 10.1038/s41593-018-0090-8.

[13] L.G. Coleman, Jr., J. Zou, F.T. Crews, Microglial depletion and repopulation in brain slice culture normalizes sensitized proinflammatory signaling, J Neuroinflammation. 17(1) (2020) 27. doi: 10.1186/s12974-019-1678-y.

[14] Y. Li, R.M. Ritzel, N. Khan, T. Cao, J. He, Z. Lei, J.J. Matyas, B. Sabirzhanov, S. Liu, H. Li, B.A. Stoica, D.J. Loane, A.I. Faden, J. Wu, Delayed microglial depletion after spinal cord injury reduces chronic inflammation and neurodegeneration in the brain and improves neurological recovery in male mice, Theranostics. 10(25) (2020) 11376–11403. doi: 10.7150/thno.49199.

[15] J.C. Perez, G. Poulen, M. Cardoso, H. Boukhaddaoui, C.M. Gazard, G. Courtand, S.S. Bertrand, Y.N. Gerber, F.E. Perrin, CSF1R inhibition at chronic stage after spinal cord injury modulates microglia proliferation, Glia. 71(12) (2023) 2782–2798. doi: 10.1002/glia.24451.

[16] W. Shi, J. Zhang, Z. Shang, Y. Zhang, Y. Xia, H. Fu, T. Yu, Restorative therapy using microglial depletion and repopulation for central nervous system injuries and diseases, Front Immunol. 13 (2022) 969127. doi: 10.3389/fimmu.2022.969127.

[17] A.K. Okojie, J.O. Uweru, M.A. Coburn, S. Li, V.D. Cao-Dao, U.B. Eyo, Distinguishing the effects of systemic CSF1R inhibition by PLX3397 on microglia and peripheral immune cells, J Neuroinflammation. 20(1) (2023) 242. doi: 10.1186/s12974-023-02924-5.

[18] Y. Liu, K.S. Given, E.L. Dickson, G.P. Owens, W.B. Macklin, J.L. Bennett, Concentration-dependent effects of CSF1R inhibitors on oligodendrocyte progenitor cells ex vivo and in vivo, Exp Neurol. 318 (2019) 32–41. doi: 10.1016/j.expneurol.2019.04.011.

[19] M.C. Operti, Y. Dölen, J. Keulen, E.A.W. van Dinther, C.G. Figdor, O. Tagit, Microfluidics-Assisted Size Tuning and Biological Evaluation of PLGA Particles, Pharmaceutics. 11(11) (2019) 590. doi: 10.3390/pharmaceutics11110590.

[20] Y. Labrak, B. Heurtault, B. Frisch, P. Saulnier, E. Lepeltier, V.E. Miron, G.G. Muccioli, A. des Rieux, Impact of anti-PDGFRα antibody surface functionalization on LNC uptake by oligodendrocyte progenitor cells, Int J Pharm. 618 (2022) 121623. doi: 10.1016/j.ijpharm.2022.121623.

[21] A. Mwema, P. Bottemanne, A. Paquot, B. Ucakar, K. Vanvarenberg, M. Alhouayek, G.G. Muccioli, A. des Rieux, Lipid nanocapsules for the nose-to-brain delivery of the anti-inflammatory bioactive lipid PGD_2_-G, Nanomedicine. 48 (2023) 102633. doi: 10.1016/j.nano.2022.

[22] C. Njoo, C. Heinl, R. Kuner, In vivo SiRNA transfection and gene knockdown in spinal cord via rapid noninvasive lumbar intrathecal injections in mice, J Vis Exp. 85 (2014) 51229. doi: 10.3791/51229.

[23] P. De Berdt, K. Vanvarenberg, B. Ucakar, C. Bouzin, A. Paquot, V. Gratpain, A. Loriot, V. Payen, B. Bearzatto, G.G. Muccioli, L. Gatto, A. Diogenes, A. des Rieux, The human dental apical papilla promotes spinal cord repair through a paracrine mechanism, Cell Mol Life Sci. 79(5) (2022) 252. doi: 10.1007/s00018-022-04210-8.

[24] A.J. Riquier, S.I. Sollars, Astrocytic response to neural injury is larger during development than in adulthood and is not predicated upon the presence of microglia, Brain Behav Immun Health. 1 (2019) 100010. doi: 10.1016/j.bbih.2019.100010.

[25] L. Cao, E.N. Bean, J.T. Malon, Preparation of Primary Mixed Glial Cell Cultures from Adult Mouse Spinal Cord Tissue, Curr Protoc. 3(4) (2023) e743. doi: 10.1002/cpz1.743.

[26] Y. Wang, I. Wernersbach, J. Strehle, S. Li, D. Appel, M. Klein, K. Ritter, R. Hummel, I. Tegeder, M.K.E. Schäfer, Early posttraumatic CSF1R inhibition via PLX3397 leads to time- and sex-dependent effects on inflammation and neuronal maintenance after traumatic brain injury in mice, Brain Behav Immun. 106 (2022) 49–66. doi: 10.1016/j.bbi.2022.07.164.

[27] A.G. Spiteri, D. Ni, Z.L. Ling, L. Macia, I.L. Campbell, M.J. Hofer, N.J.C. King, PLX5622 Reduces Disease Severity in Lethal CNS Infection by Off-Target Inhibition of Peripheral Inflammatory Monocyte Production, Front Immunol. 13 (2022) 851556. doi: 10.3389/fimmu.2022.851556.

[28] F. Lei, N. Cui, C. Zhou, J. Chodosh, D.G. Vavvas, E.I. Paschalis, CSF1R inhibition by a small-molecule inhibitor is not microglia specific; affecting hematopoiesis and the function of macrophages, Proc Natl Acad Sci U S A. 117(38) (2020) 23336–23338. doi: 10.1073/pnas.1922788117.

[29] M.G. Lampugnani, M. Pedenovi, A. Niewiarowski, B. Casali, M.B. Donati, G.C. Corbascio, P.C. Marchisio, Effects of dimethyl sulfoxide (DMSO) on microfilament organization, cellular adhesion, and growth of cultured mouse B16 melanoma cells, Exp Cell Res. 172(2) (1987) 385–96. doi: 10.1016/0014-4827(87)90396-x.

[30] A.F. Davidson, C. Glasscock, D.R. McClanahan, J.D. Benson, A.Z. Higgins, Toxicity Minimized Cryoprotectant Addition and Removal Procedures for Adherent Endothelial Cells, PLoS One. 10(11) (2015) e0142828. doi: 10.1371/journal.pone.0142828.

[31] M. Awan, I. Buriak, R. Fleck, B. Fuller, A. Goltsev, J. Kerby, M. Lowdell, P. Mericka, A. Petrenko, Y. Petrenko, O. Rogulska, A. Stolzing, G.N. Stacey, Dimethyl sulfoxide: a central player since the dawn of cryobiology, is efficacy balanced by toxicity? Regen Med. 15(3) (2020) 1463–1491. doi: 10.2217/rme-2019-0145.

[32] S.J. Shepherd, X. Han, A.J. Mukalel, R. El-Mayta, A.S. Thatte, J. Wu, M.S. Padilla, M.G. Alameh, N. Srikumar, D. Lee, D. Weissman, D. Issadore, M.J. Mitchell, Throughput-scalable manufacturing of SARS-CoV-2 mRNA lipid nanoparticle vaccines, Proc Natl Acad Sci U S A. 120(33) (2023) e2303567120. doi: 10.1073/pnas.2303567120.

[33] B. Hu, S. Duan, Z. Wang, X. Li, Y. Zhou, X. Zhang, Y.W. Zhang, H. Xu, H. Zheng, Insights Into the Role of CSF1R in the Central Nervous System and Neurological Disorders, Front Aging Neurosci. 13 (2021) 789834. doi: 10.3389/fnagi.2021.789834.

[34] D.P. Sester, A. Trieu, K. Brion, K. Schroder, T. Ravasi, J.A. Robinson, R.C. McDonald, V. Ripoll, C.A. Wells, H. Suzuki, Y. Hayashizaki, K.J. Stacey, D.A. Hume, M.J. Sweet, LPS regulates a set of genes in primary murine macrophages by antagonising CSF-1 action, Immunobiology. 210(2-4) (2005) 97–107. doi: 10.1016/j.imbio.2005.05.004.

[35] E. Rovida, A. Paccagnini, M. Del Rosso, J. Peschon, P. Dello Sbarba, TNF-alpha-converting enzyme cleaves the macrophage colony-stimulating factor receptor in macrophages undergoing activation, J Immunol. 166(3) (2001) 1583–9. doi: 10.4049/jimmunol.166.3.1583.

[36] X. Du, Y. Xu, S. Chen, M. Fang, Inhibited CSF1R Alleviates Ischemia Injury via Inhibition of Microglia M1 Polarization and NLRP3 Pathway, Neural Plast. 2020 (2020) 8825954. doi: 10.1155/2020/8825954.

[37] S. Lee, X.Q. Shi, A. Fan, B. West, J. Zhang, Targeting macrophage and microglia activation with colony stimulating factor 1 receptor inhibitor is an effective strategy to treat injury-triggered neuropathic pain, Mol Pain. 14 (2018) 1744806918764979. doi: 10.1177/1744806918764979.

[38] E.F. Willis, K.P.A. MacDonald, Q.H. Nguyen, A.L. Garrido, E.R. Gillespie, S.B.R. Harley, P.F. Bartlett, W.A. Schroder, A.G. Yates, D.C. Anthony, S. Rose-John, M.J. Ruitenberg, J. Vukovic, Repopulating Microglia Promote Brain Repair in an IL-6-Dependent Manner, Cell. 180(5) (2020) 833–846.e16. doi: 10.1016/j.cell.2020.02.013.

[39] E.G. Vichaya, S. Malik, L. Sominsky, B.G. Ford, S.J. Spencer, R. Dantzer, Microglia depletion fails to abrogate inflammation-induced sickness in mice and rats, J Neuroinflammation. 17(1) (2020) 172. doi: 10.1186/s12974-020-01832-2.

[40] T. Clifford, Z. Finkel, B. Rodriguez, A. Joseph, L. Cai, Current Advancements in Spinal Cord Injury Research-Glial Scar Formation and Neural Regeneration, Cells. 12(6) (2023) 853. doi: 10.3390/cells12060853.

[41] W. Luo, Y. Li, C. Xiang, T. Aizawa, R. Niu, Y. Wang, J. Zhao, Z. Liu, C. Li, W. Liu, R. Gu, Nanomaterials as therapeutic agents to modulate astrocyte-mediated inflammation in spinal cord injury, Mater Today Bio. 23 (2023) 100888. doi: 10.1016/j.mtbio.2023.100888.

[42] K.G. Witcher, C.E. Bray, T. Chunchai, F. Zhao, S.M. O’Neil, A.J. Gordillo, W.A. Campbell, D.B. McKim, X. Liu, J.E. Dziabis, N. Quan, D.S. Eiferman, A.J. Fischer, O.N. Kokiko-Cochran, C. Askwith, J.P. Godbout, Traumatic Brain Injury Causes Chronic Cortical Inflammation and Neuronal Dysfunction Mediated by Microglia, J Neurosci. 41(7) (2021) 1597–1616. doi: 10.1523/JNEUROSCI.2469-20.2020.

[43] D.E. Marzan, V. Brügger-Verdon, B.L. West, S. Liddelow, J. Samanta, J.L. Salzer, Activated microglia drive demyelination via CSF1R signaling, Glia. 69(6) (2021) 1583–1604. doi: 10.1002/glia.23980.

[44] B. Basilico, L. Ferrucci, A. Khan, S. Di Angelantonio, D. Ragozzino, I. Reverte, What microglia depletion approaches tell us about the role of microglia on synaptic function and behavior, Front Cell Neurosci. 16 (2022) 1022431. doi: 10.3389/fncel.2022.1022431.

[45] H. Fu, Y. Zhao, D. Hu, S. Wang, T. Yu, and L. Zhang, Depletion of microglia exacerbates injury and impairs function recovery after spinal cord injury in mice, Cell Death Dis. 11(7) (2020) 528. doi: 10.1038/s41419-020-2733-4.

[46] G.J. Duncan, S.B. Manesh, B.J. Hilton, P. Assinck, J.R. Plemel, W. Tetzlaff, The fate and function of oligodendrocyte progenitor cells after traumatic spinal cord injury, Glia. 68(2) (2020) 227–245. doi: 10.1002/glia.23706.

[47] I. Francos-Quijorna, J. Amo-Aparicio, A. Martinez-Muriana, R. López-Vales, IL-4 drives microglia and macrophages toward a phenotype conducive for tissue repair and functional recovery after spinal cord injury, Glia. 64(12) (2016) 2079–2092. doi: 10.1002/glia.23041.

[48] X.J. Wang, G.F. Shu, X.L. Xu, C.H. Peng, C.Y. Lu, X.Y. Cheng, X.C. Luo, J. Li, J. Qi, X.Q. Kang, F.Y. Jin, M.J. Chen, X.Y. Ying, J. You, Y.Z. Du, J.S. Ji, Combinational protective therapy for spinal cord injury medicated by sialic acid-driven and polyethylene glycol based micelles, Biomaterials. 217 (2019) 119326. doi: 10.1016/j.biomaterials.2019.119326.

