## Supplementary Data for "Nanoparticles loaded with a CSF1R antagonist selectively depletes microglial cells and modulates inflammation in spinal cord injury"

### Tables

**Table S1.** Antibodies used for immunofluorescence and FACS.

| Primary Antibody | Ref | Supplier | Dilution |
|---|---|---|---|
| Anti- Iba1 | 019-19741 | Wako (Sopachem) | 1:1000 |
| Anti- GFAP | ab7779 | Abcam | 1:1000 |
| Anti- olig2 | P21954 | Invitrogen (Thermo) | 1:1000 |
| Anti- olig2 | ab109186 | Abcam | 1:500 |
| Anti- NeuN | MAB377 | Millipore | 1:1000 |
| Anti- CD86 | 105023 | Biolegend | 1:200(IF)/ 1:100(FACS) |
| Anti- CD206 | 141711 | Biolegend | 1:200(IF)/ 1:100(FACS) |
| Anti- F4/80 | 565854 | BD Pharmingen™ | 1:100(FACS) |

| Secondary antibody | Ref | Supplier | Dilution |
|---|---|---|---|
| Alexa 488 | A11008 | Invitrogen (Thermo) | 1:500 |
| Alexa 594 | A21125 | Invitrogen (Thermo) | 1:500 |
| Alexa 647 | A21247 | Invitrogen (Thermo) | 1:500 |

**Table S2.** Primers used for qRT-PCR.

Mouse

| Target | Forward (5'-3') | Reverse (5'-3') |
|---|---|---|
| Arg-1 | GGTTCTGGGAGGCCTATCTT | TGAAAGGAGCCCTGTCTTGT |
| CD11b | GCTGGGGGACAGTAGAAACA | GCTGGGGGACAGTAGAAACA |
| CD206 | CCGTCTGTGCATTTCCATTC | TGCCAGTCAGTGGATCTTTGT |
| CD86 | TTGTGTGTGTTCTGGAAACGGAG | AACTTAGAGGCTGTGTTGCTGGG |
| CSF1R | ACAAGTACAAGCAGAAGCCGA | GCACTGCATCTTCTTTGCCC |
| Fizz | AAGGAACTTCTTGCCAATCCA | CACAAGCACACCCAGTAGCA |
| GFAP | GCTGGGCCTAGGGGATTATG | AACGCATCTCTCCATCGCTT |
| Iba-1 | CCTGAGGAGATTTCAAAAGCTGA | TGGGACCGTTCTCACACTTC |
| IL-1β | TCGCTCAGGGTCACAAGAAA | CATCAGAGGCAAGGAGGAAAAC |
| IL-33 | GCCTCCCTGAGTACATACAATG | CGTAGTAGCACCTGGTCTTG |
| IL-6 | ACAAGTCGGAGGCTTAATTACACAT | TTGCCATTGCACAACTCTTTTC |
| iNOS | AGGTACTCAGCGTGCTCCAC | GCACCGAAGATATCTTCATG |
| Olig2 | AACCGCATCACCATTCCTGT | CAGGCTGGTTTCTCGGATCT |
| RPL19 | TGACCTGGATGAGAAGGATGAG | CTGTGATACATATGGCGGTCAATC |

| | | |
|---|---|---|
| TNF-α | AGCCCCCAGTCTGTATCCTT | GGTCACTGTCCCAGCATCTT |
| Ym-1 | GCTTTTGAGGAAGAATCTGTGGA | AAGAGACTGAGACAGTTCAGGGAT |

Rat

| Target | Forward (5'-3') | Reverse (5'-3') |
|---|---|---|
| Arg-1 | GAAGGTCTCTACATCACAGAAGAAA | CAAGGTCAACGCCACTGC |
| CSF1R | GGACCACATACAGCTACTCATTC | AGATTATTCCAGCCTGCCTTG |
| IL-10 | CGACGCTGTCATCGATTTCTC | CAGTAGATGCCGGGTGGTTC |
| iNOS | TCACCTATCGCACCCGAG | AAGCCACTGACACTCCGC |
| RPL13 | GGCTGAAGCCTACCAGAAAG | CTTTGCCTTTTCCTTCCGTT |
| TNF-α | AGTGACAAGCCCGTAGCC | TTGAAGAGAACCTGGGAGTAGA |

### Figures

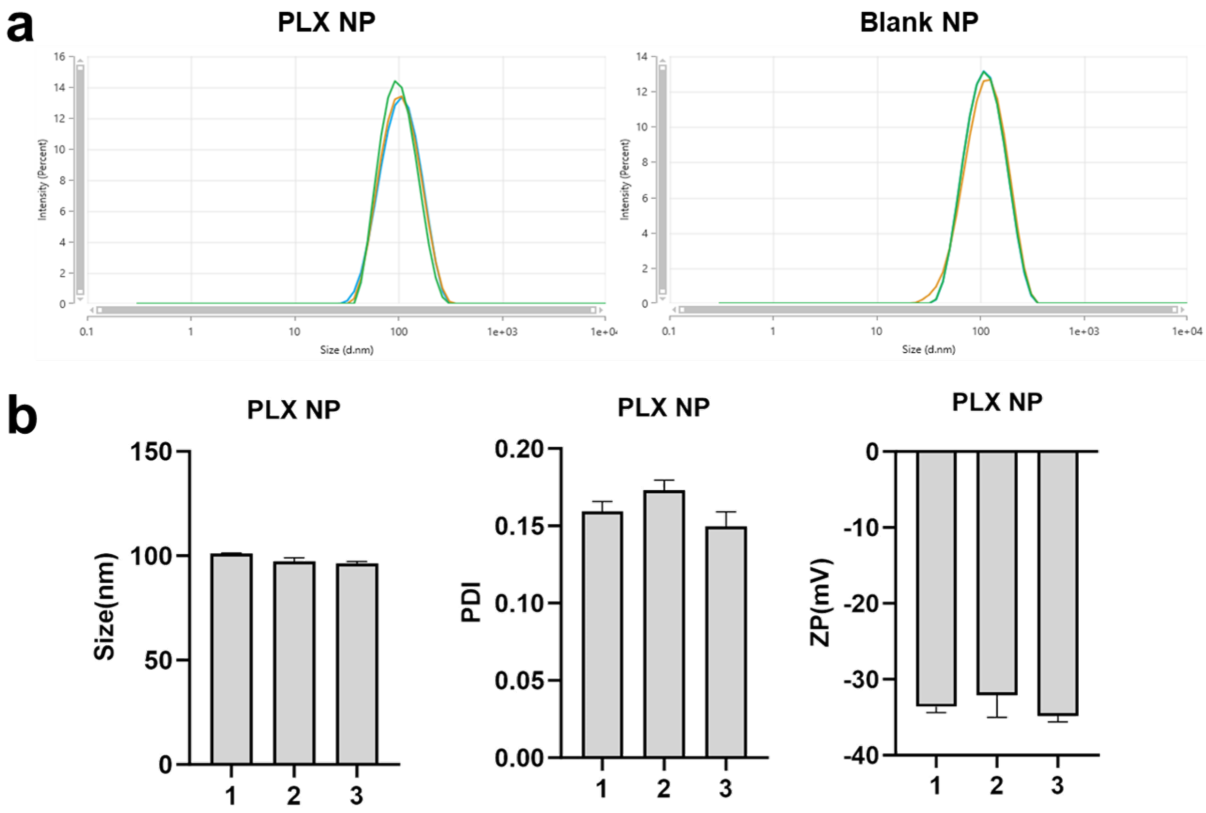


c

| | Size (nm) | PDI | Zeta potential(mV) |
|---|---|---|---|
| PLX NP- DiD | 152± 2 | 0.15± 0.03 | -27 ± 0.5 |

**Figure S1.** Nanoparticle characterization. PLX NP and Blank NP size graph (a) were measured by nanosizer. (b) PLX NP size, PDI and Zeta potential for 3 batches of the same formulation. (c) Physicochemical characterization of PLX NP- DiD.

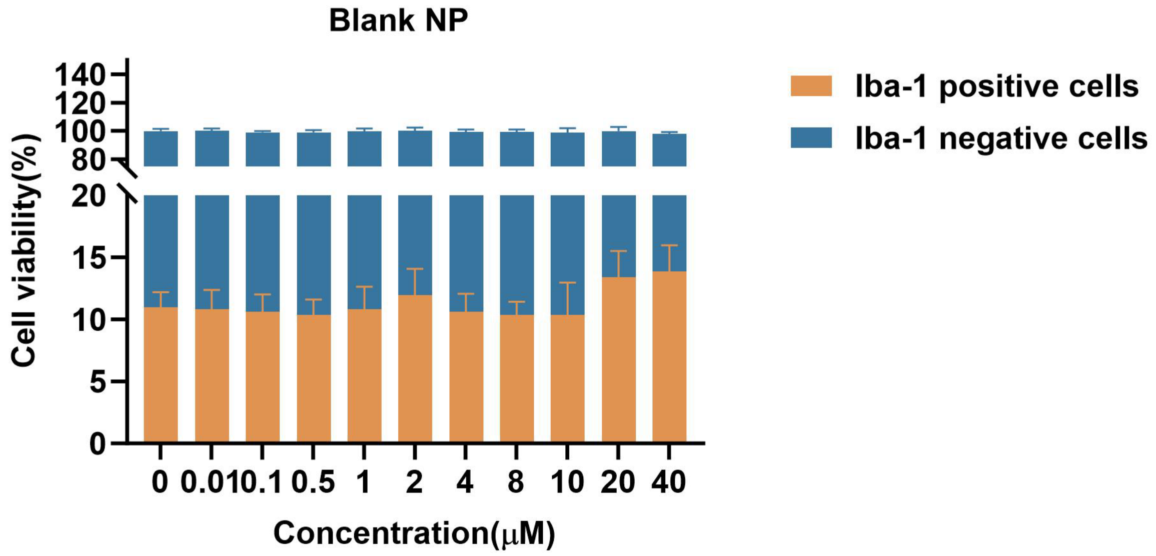


**Figure S2.** Impact of Blank NP on selective depletion of microglial cells in vitro. MGC cells were incubated with different concentrations of Blank NP for 72 h (d) (N=3, n=8). Then cells were stained for Iba-1 (microglial cells) and DAPI (nuclei) and images were acquired with an IX-PICO. Analysis was done on whole wells using IX-PICO software. Data are expressed as the mean ± SD. Statistical analysis was done using a Nested one-way ANOVA or two-way ANOVA (*$p < 0.05$).

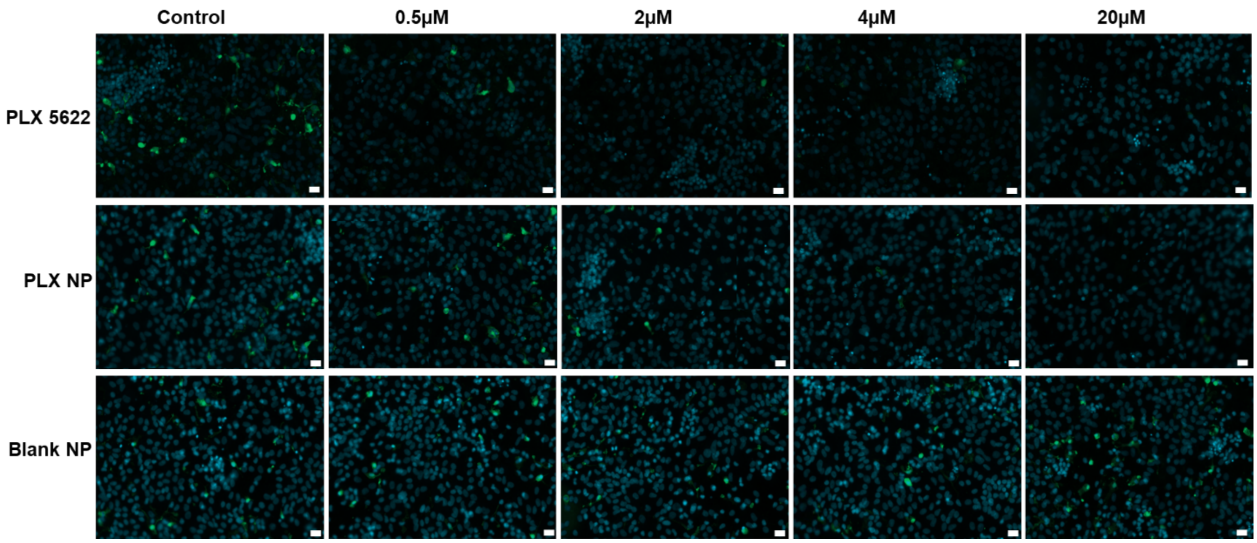


**Figure S3.** Impact of PLX NP on selective depletion of microglial cells in vitro. MGC cells were incubated with different concentrations of PLX, PLX NP, and Blank NP for 72h (N=3, n=8). Then cells were stained for Iba-1 (microglial cells) and DAPI (nuclei) and images were acquired with an IX-PICO. Scale bar: 20 µm, DAPI-blue, Iba1-green.

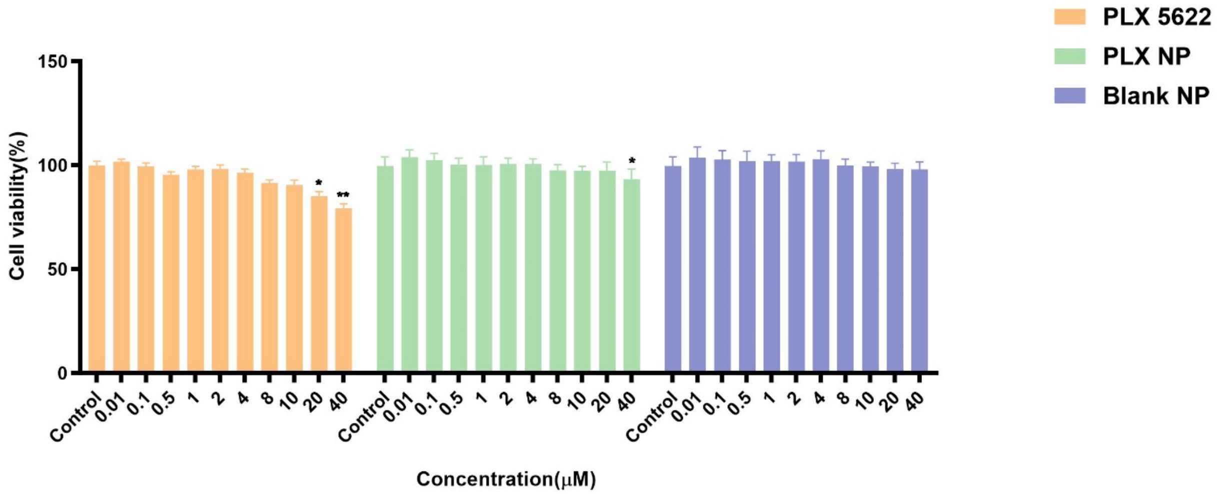


**Figure S4.** PLX NP cytotoxicity on MGC. MGC cells were incubated with different concentrations of PLX, PLX NP, and Blank NP for 72h (N=3, n=8). Then cell viability was tested by Presto Blue and acquired with Spectramax. Data are expressed as the mean ± SD. Statistical analysis was done using a Nested one-way ANOVA (*$p < 0.05$).

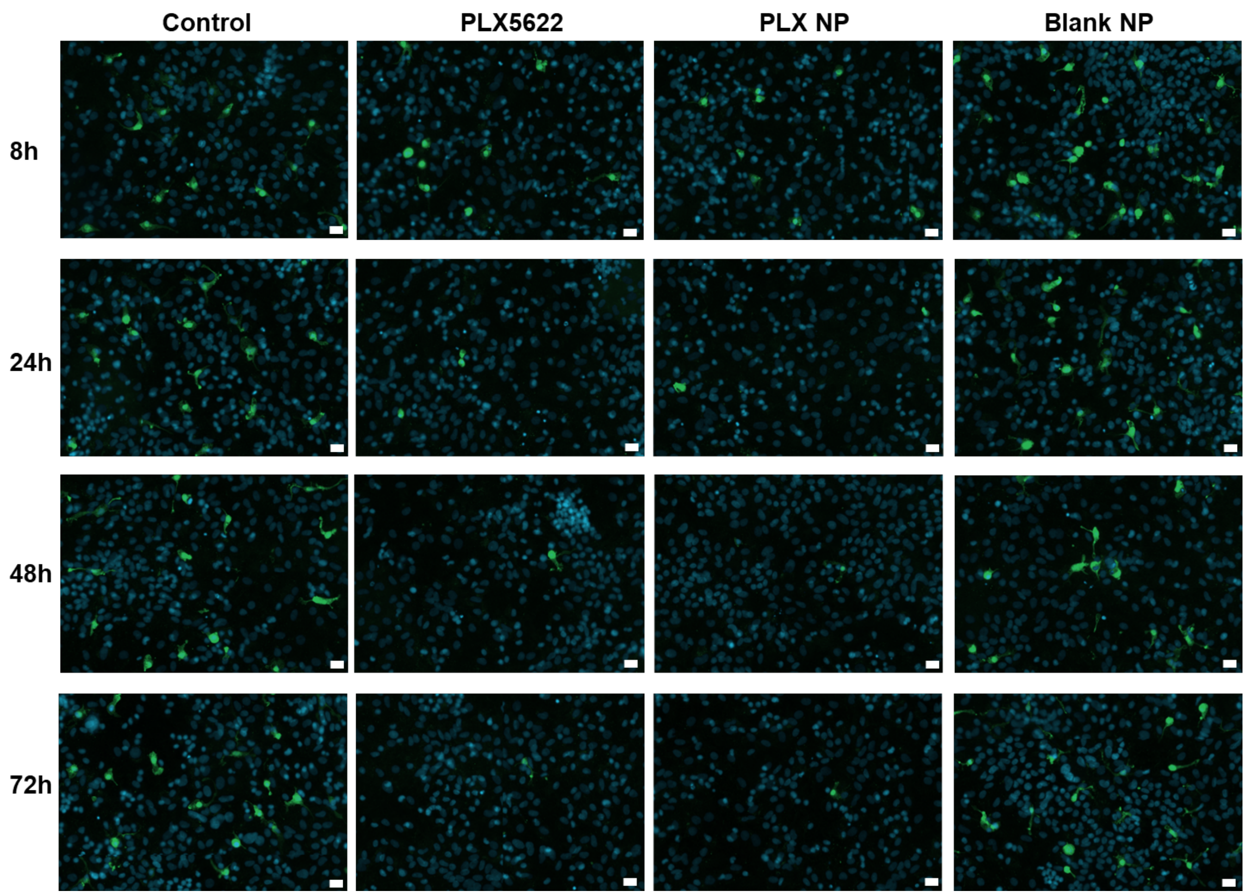


**Figure S5.** Impact of PLX NP incubation time on microglial depletion. MGC cells were incubated with 4 μM of PLX, PLX NP and Blank NP for 8 h, 24 h 48 h and 72 h (N=3, n=6). Then cells were stained for Iba-1 (microglial cells) and DAPI (nuclei) and images were acquired with an IX-PICO. Scale bar: 20 μm, DAPI-blue, Iba1-green.

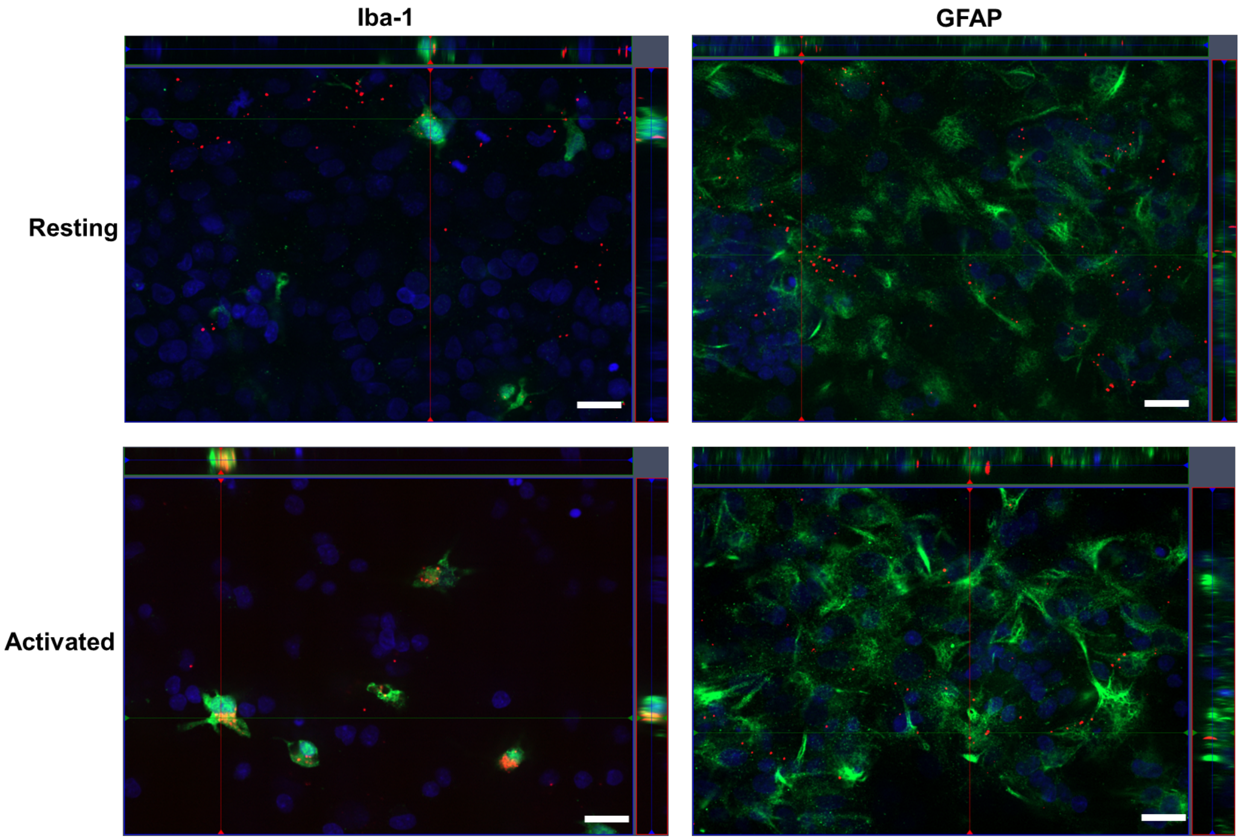


**Figure S6.** Uptake of PLX NP- DiD by MGC activated (Activated) or not (Resting) by LPS. MGC were incubated with 4 µM of PLX NP- DiD for 1 h and 24 h (N=3, n=4). Microglial cells were stained for Iba-1, astrocytes for GFAP, oligodendrocyte lineage cells for Olig2 and nuclei with DAPI. Images were acquired with an ZEISS and IX-PICO for the quantification. Figure 5a scale bar: 20 µm, DAPI- blue, Iba-1-green, GFAP- green, PLX NP- DiD- red.

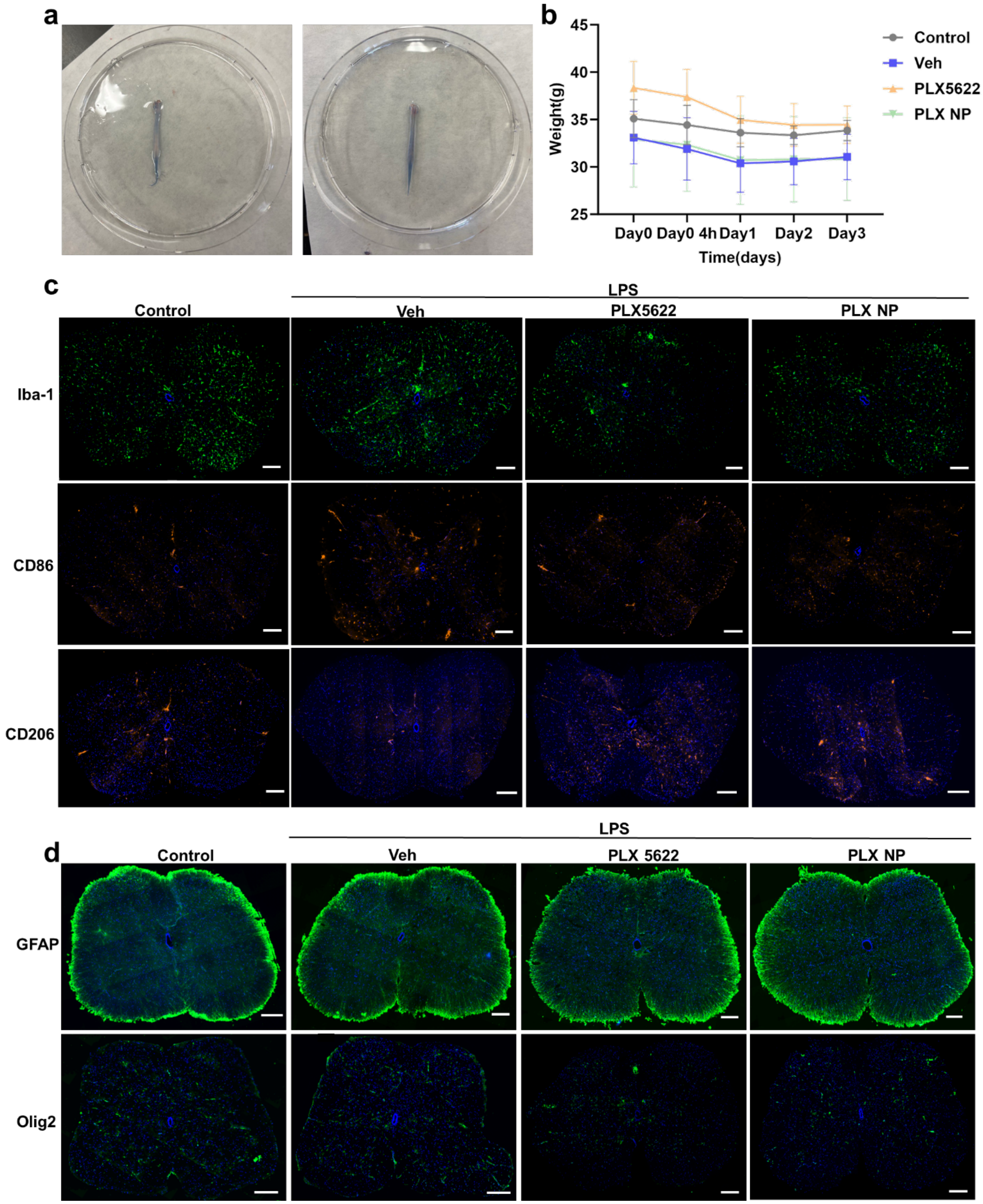


**Figure S7.** PLX NP impact on LPS-activated mice after intrathecal injection. (a) Spinal cord was dissected following intrathecal injection with blue dye. (b) Weight over time (g); Spinal cords were immunostained for (b) Iba-1 (microglia cells), CD86 (M1-like phenotype microglia), CD206 (M2-

like phenotype microglia), (c) GFAP (astrocytes), Olig2 (oligodendrocyte lineage) and DAPI (nuclei) and images were acquired with a slide scanner. Scale bar: 200 µm, DAPI- blue, Iba-1- green, CD86-red, CD206-red, GFAP- green, Olig2- green.

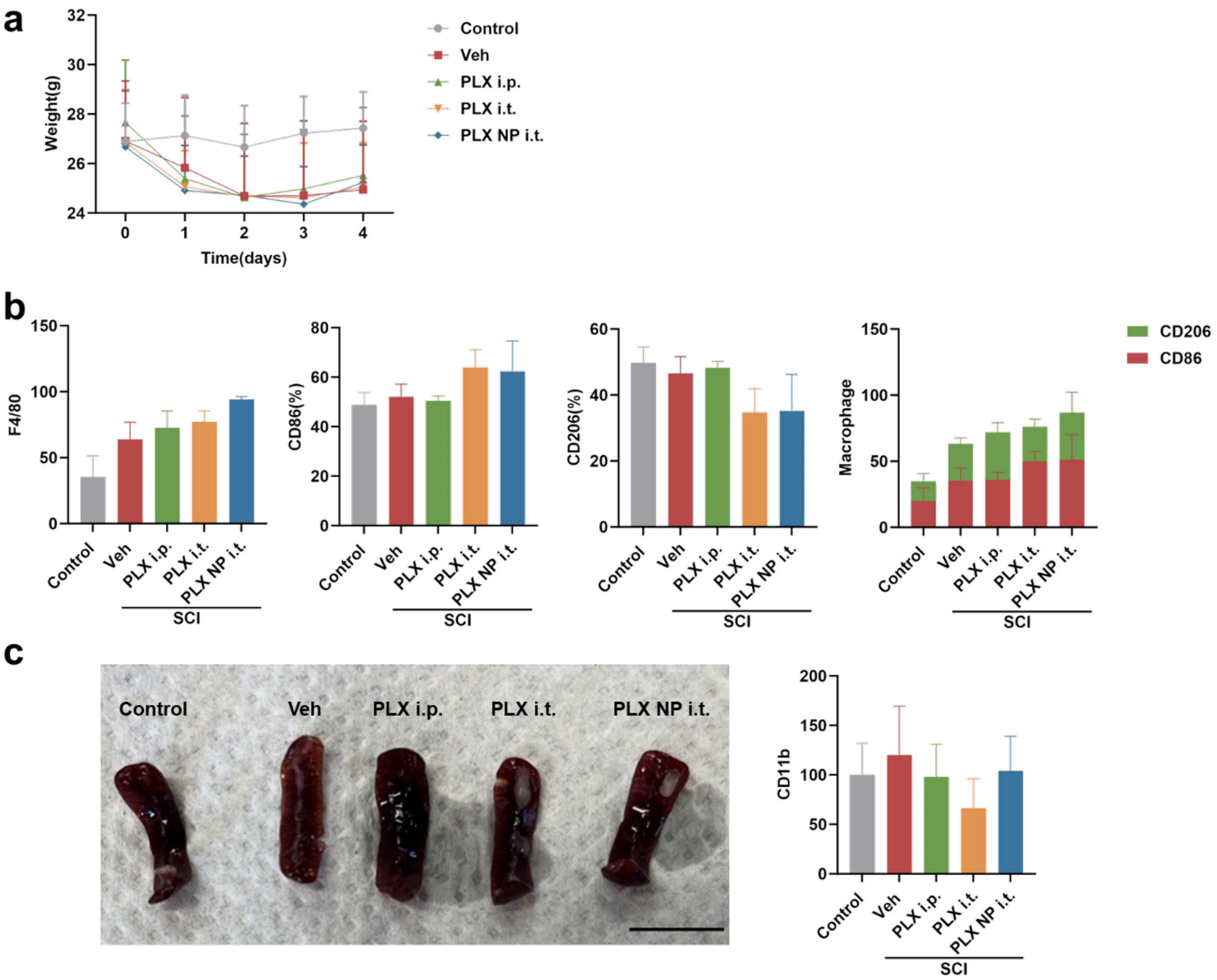


**Figure S8.** Impact of PLX NP on circulating macrophages in a mouse contusion SCI model. (a) Weight over time (g); (b) Whole blood FACS for F4/80 (macrophage), CD86 (M1-like macrophage), and CD206 (M2-like macrophage); (c) Spleen macroscopic appearance CD11b gene expression. Scale bar: 1 cm. Data are expressed as the mean ± SEM. Statistical analysis was done using One-way ANOVA.

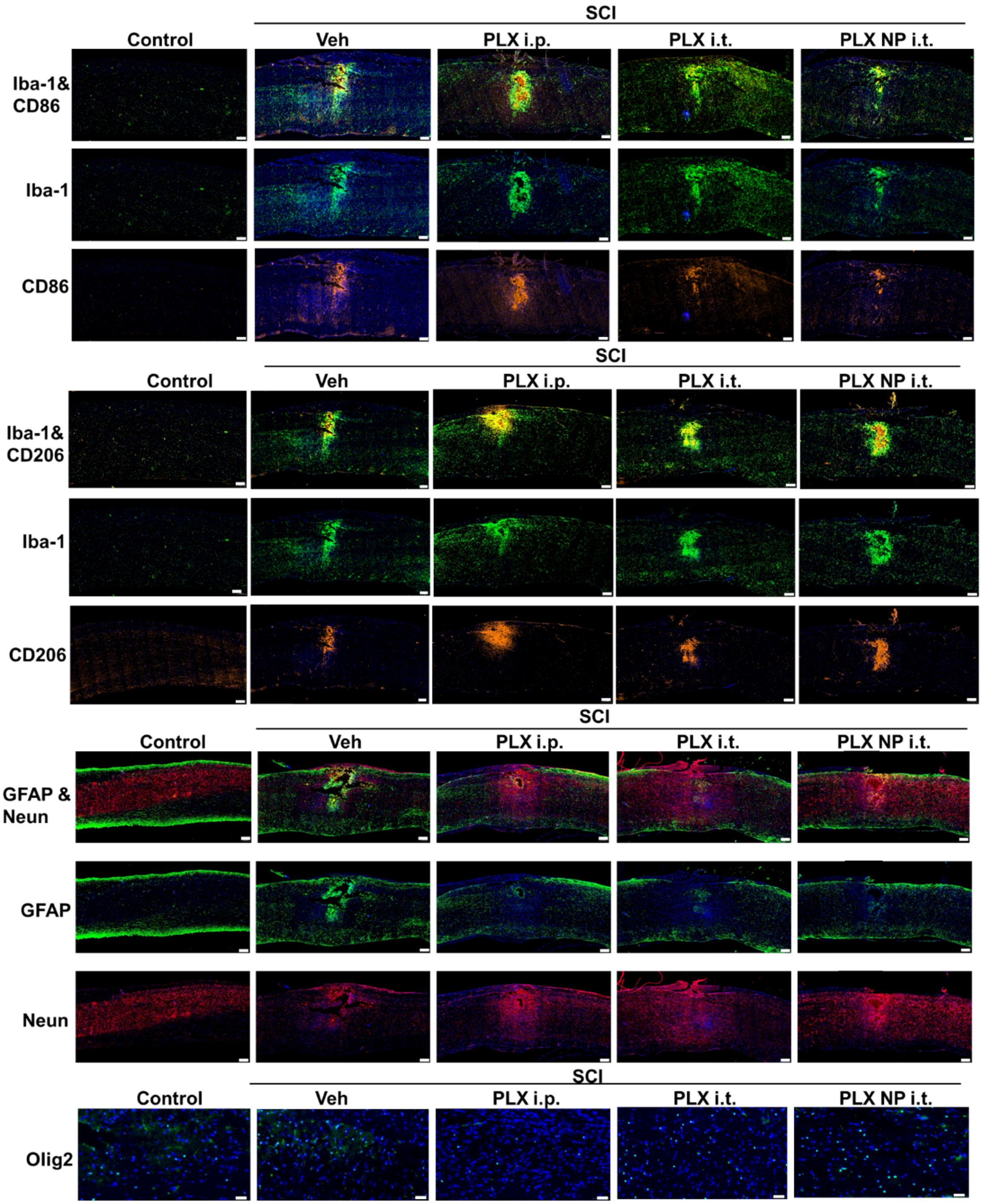


**Figure S9.** PLX NP effect on SCI microenvironment. Spinal cords were immunostained for Iba-1 (microglia cells), CD86 (M1-like phenotype microglia), CD206 (M2-like phenotype microglia), GFAP (astrocytes), Olig2 (oligodendrocyte lineage) and DAPI (nuclei) and images were acquired with a slide scanner. Iba-1, CD86, CD206, GFAP and Neun scale bar: 200 µm, Olig2 scale bar: 50 µm. DAPI- blue, Iba-1-green, CD86-red, CD206-red, GFAP- green, Neun- red, Olig2- green.

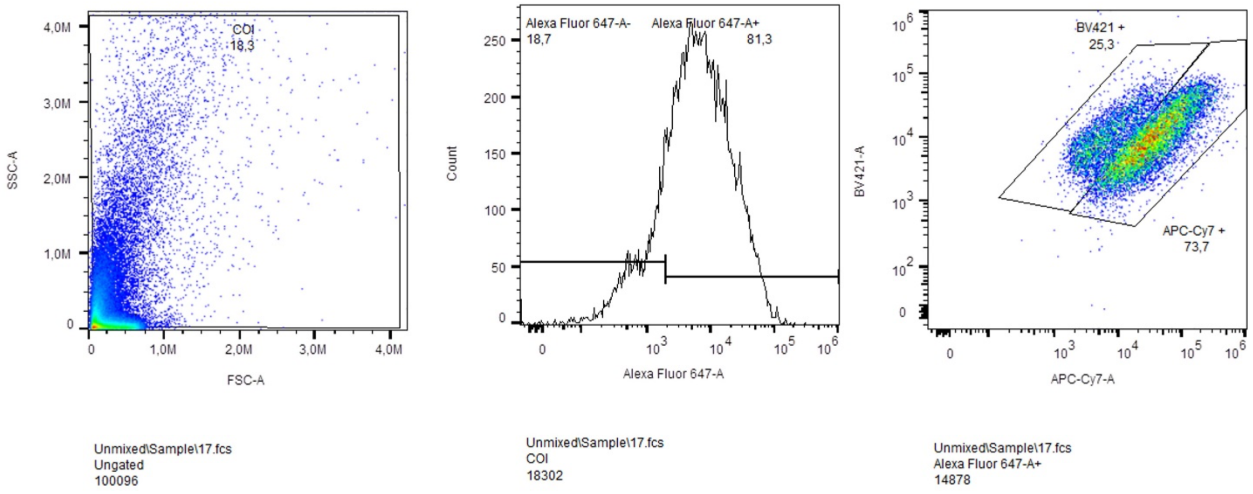


**Figure S10.** Settings used to analyze the whole blood of mice by FACS.
